# Germline-specific role for unconventional components of the γ-tubulin complex in *Caenorhabditis elegans*

**DOI:** 10.1101/2022.12.23.521729

**Authors:** Nami Haruta, Eisuke Sumiyoshi, Yu Honda, Masahiro Terasawa, Chihiro Uchiyama, Mika Toya, Yukihiko Kubota, Asako Sugimoto

**Author notes:** Institute of Molecular Biology, University of Oregon, Eugene, OR 97403. Department of Life Science and Medical Bioscience, School of Advanced Science and Engineering, Waseda University, Shinjuku-ku, Tokyo 162-8480, Japan (Mika Toya). Laboratory of Information Biology, College of life Sciences, Department of Bioinformatics, Ritsumeikan University, Kusatsu, Shiga 525-8577, Japan (Yukihiko Kubota). Corresponding authors: Laboratory of Developmental Dynamics, Graduate School of Life Sciences, Tohoku University, 2-1-1 Katahira, Aoba-ku, Sendai 980-8577, Japan.

## Abstract

The γ-tubulin complex (γTuC) is a widely conserved microtubule nucleator, but some of its components GCP4–6. Here, we identified two γTuC-associated proteins in *C. elegans*, namely GTAP−1 and −2, for which apparent orthologs were detected only in the genus *Caenorhabditi*s. Their centrosomal localization was interdependent. In early *C. elegans* embryos, whereas the conserved γTuC component MZT-1/MOZART1 was essential for the localization of centrosomal γ-tubulin, depletion of GTAP-1 and/or −2 caused up to 50% reduction of centrosomal γ-tubulin and precocious disassembly of spindle poles during mitotic telophase. In the adult germline, GTAP-1 and GTP-2 contributed to the efficient recruitment of γTuC to the plasma membrane. Depletion of GTAP-1, but not GTAP-2, severely disrupted both the microtubule array and the honeycomb-like structure in the adult germline.

We propose that GTAP-1 and −2 are unconventional components of γTuC that contribute to the organization of both centrosomal and non-centrosomal microtubules by targeting the γTuC to specific subcellular sites in a tissue-specific manner.

**SUMMARY STATEMENT:** Haruta et al. show that GTAP-1 and −2, two unconventional components of the γ-tubulin complex (γTuC) in *C. elegans*, contribute to targeting the γTuC to embryonic centrosomes and germ-cell membrane in adults.

## INTRODUCTION

The γ-tubulin complex (γTuC) is a highly conserved microtubule (MT) nucleator that accumulates at microtubule-organizing centers (MTOCs) such as centrosomes in animal cells and spindle-pole bodies in yeasts. There are two types of γTuCs with different sizes: the γ-tubulin small complex (γTuSC) and the γ-tubulin ring complex (γTuRC) (Kollman et al., 2011). The γTuSC consists of two molecules of γ-tubulin and one molecule each of GCP2 and GCP3; two γ-tubulin molecules bind to the C-terminal region of GCP2 and GCP3, forming a Y-shaped structure (Guillet et al., 2011; Kollman et al., 2008). The γTuRC comprises four to six γTuSCs and several additional γTuRC-specific components including GCP4–6 and the small protein MOZART1. The γTuRC forms a lock-washer-shaped structure to provide a template for 13-protofilament MTs (Zheng et al., 1995; Moritz et al., 1998; Oegema et al., 1999; Keating and Borisy, 2000; Wiese and Zheng, 2000; Murphy et al., 2001; Kollman et al., 2010). GCP2–6 share common structures, containing two conserved regions called grip1 and grip2 motifs; the grip2 motif directly binds γ-tubulin, whereas grip1 is involved in lateral interaction between GCPs (Gunawardane et al., 2000; Guillet et al., 2011; Kollman et al., 2008). GCP4–6 are proposed to be integrated into γTuRC as a spoke similar to γTuSC assembly at the distal seam of the ring (Liu et al., 2019; Wieczorek et al., 2020; Consolati, et al., 2020).

Among the γTuC components, the requirement of GCP4–6 for γTuC function varies depending on the tissue and organism (Anders et al., 2006; Colombie et al., 2006; Venkatram et al., 2004; Fujita et al., 2002). In particular, *Saccharomyces cerevisiae* lacks GCP4–6 (Teixido-Travesa et al., 2012; Kollman et al., 2010; Geissler et al., 1996; Knop and Schiebel, 1997), but this yeast’s γTuSCs can form a stable 13-symmetry ring-like structure by associating with Spc110 (Kollman et al., 2010; Kollman et al., 2015). In *Drosophila*, Grip75/GCP4 and Grip128/GCP5, which correspond to γTuRC-specific components, are dispensable for viability but required for spermatogenesis (Vogt et al., 2006). Thus, GCP4–6 functions have diverged during evolution to play various roles in different cell types.

The nematode *Caenorhabditis elegans* is another organism in which the genes encoding GCP4–6 have not been identified in the genome. The *C. elegans* gene *tbg-1* is the only γ-tubulin gene (Bobinnec et al., 2000; Strome et al., 2001; Hannak et al., 2002), and GIP-2/GCP2, GIP-1/GCP3 and MZT-1/MOZART1 were identified based on their sequence similarities (Hannak et al., 2002; Sallee et al., 2018). Whether the *C. elegans* γTuC contains additional components remains unknown.

The γTuC in *C. elegans* localizes to various sites in a cell type–specific or tissue-specific manner. In mitotic cells, centrosomal localization of γTuC is dependent on SPD-5, a scaffold protein of the pericentriolar matrix (PCM) and a functional homolog of *Drosophila* Centrosomin (Hamill, et al., 2002; Nakajo, et al., 2022). In some tissues, the γTuC localizes at non-centrosomal MTOCs to organize specific MT arrays (Bobinnec et al., 2000; Zhou et al., 2009; Wang et al., 2015; Sallee et al., 2018; Sanchez et al., 2021). For example, in embryonic intestinal cells, γTuC localizes to the apical membrane with the tissue-specific protein WDR-62 but does not require MZT-1 (Sallee et al., 2018; Sanchez et al., 2021). In the adult hermaphrodite germline, the γTuC along with the NINEIN homolog NOCA-1 localizes at the germ-cell membrane, and this localization is required to maintain the germline morphology (Green et al., 2011; Wang et al., 2015). This tissue-specific localization and function of γTuC imply that certain unidentified components or associated proteins may be involved in the targeting of γTuC to specific intracellular sites in different tissues.

Here, we report the identification and characterization of two γ-tubulin-associated proteins, namely GTAP-1 and −2, in *C. elegans*. These proteins directly interact with γTuC components and are predicted to have a partial structural similarity to the GCP components of γTuC. Both proteins contribute to the efficient recruitment of γ-tubulin to centrosomes in early embryos and the plasma membrane in the germline. GTAP-1, but not GTAP-2, plays a crucial role in the organization of non-centrosomal MTs in the germline. We propose that GTAP-1 and GTAP-2 are highly diverged GCP components of γTuC in *C. elegans* that play distinct roles in the centrosomal and non-centrosomal MTOCs in a tissue-specific manner.

## RESULTS

### Identification of novel γ-tubulin-associated proteins in *C. elegans*

To identify new components of the γTuC in *C. elegans*, we biochemically analyzed proteins that associate with γ-tubulin. An antibody specific for FLAG was used for immunoprecipitation of an extract of worm embryos expressing GFP::γ-tubulin::FLAG. The immunoprecipitates were comprehensively analyzed by mass spectrometry (Fig. 1A,B). The known components of the γTuSC, namely GIP-1/GCP3 and GIP-2/GCP2, efficiently co-precipitated with γ-tubulin, as expected (Fig. 1A,B). Two uncharacterized proteins, encoded by *ZK1248.13* and *ZK632.5*, were identified in the top-ranked list of proteins that co-precipitated with γ-tubulin; the proteins were named GTAP-1 (gamma-tubulin-associated protein 1) and GTAP-2, respectively.

**Fig. 1.**
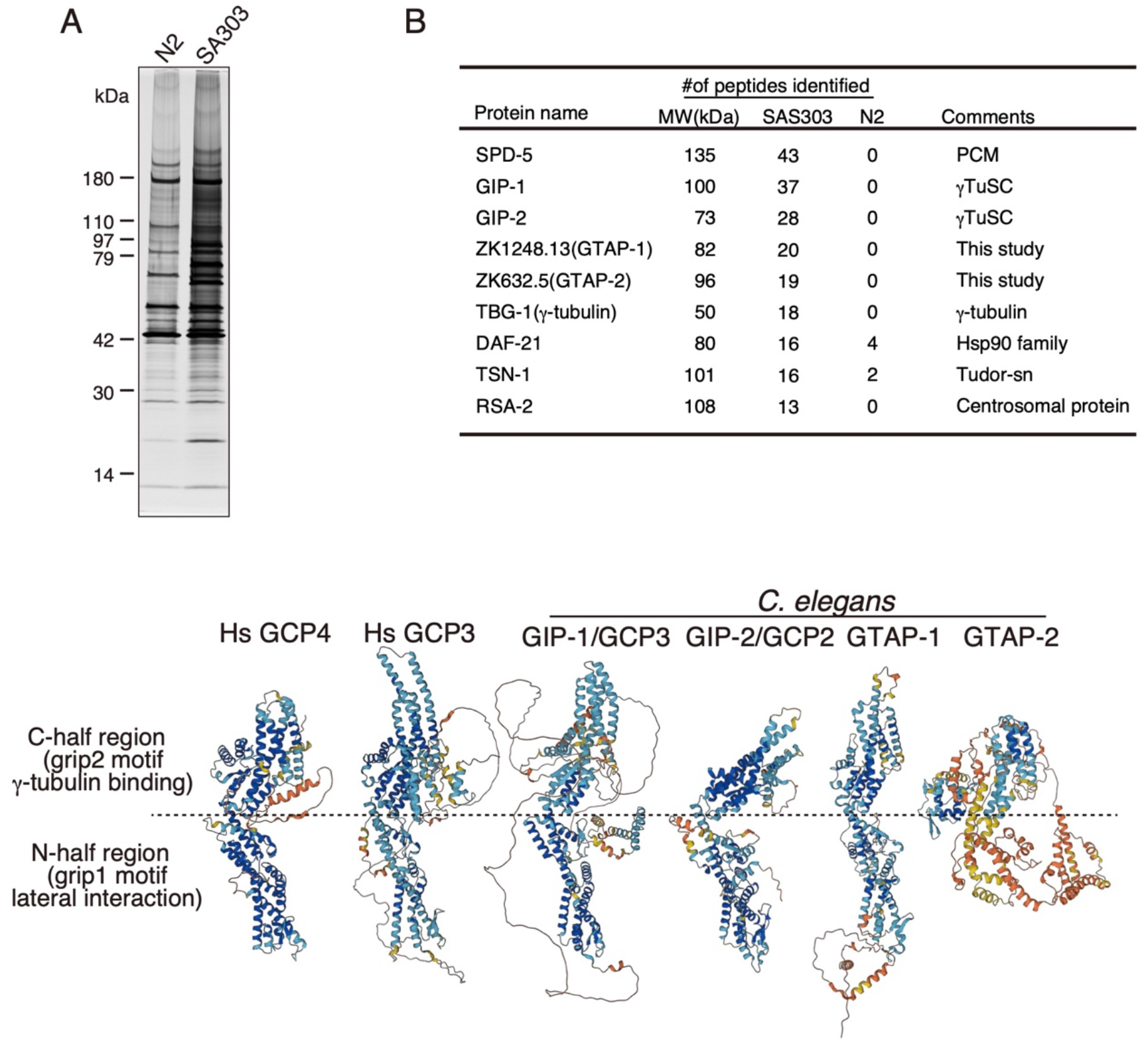
GTAP-1 and GTAP-2 are γ-tubulin-associated proteins. (A) Silver-stained SDS-PAGE gel of immunoprecipitates obtained from the control (N2) and GFP::TBG-1/γ-tubulin::FLAG-expressing strain (SA303) using anti-FLAG. The positions of identified proteins are indicated. (B) List of abundant peptides identified in the GFP:: TBG-1/γ-tubulin::FLAG precipitates. GTAP-1 (ZK1248.13) and GTAP-2 (ZK632.5) were identified together with the γTuSC components GIP-1/GCP3 and GIP-2/GCP2. (C) Predicted protein structures of human GCP4 and GCP3 as well as *C. elegans* GIP-1/GCP3, GIP-2/GCP2, GTAP-1 and GTAP-2 modeled by AlphaFold (Jumper, et.al., 2021).

GTAP-1 and −2 lacked any significant amino-acid sequence similarity to known proteins based on BLAST (Basic Local Alignment Search Tool) searches, with the exception of apparent orthologs of nematodes in the genus *Caenorhabditis*. However, the overall predicted structure of GTAP-1 modeled with AlphaFold (Jumper, et.al., 2021) resembled that of the human GCPs and *C. elegans* GIP-1/GCP3 (Fig. 1 C). A region of GTAP-2 showed only modest sequence similarity in a part of the grip-2 motif, which corresponds to a γ-tubulin-binding site conserved among GCP2–6 (Fig. S1), but the similarity with the grip-1 motif was not detected. The local sequence similarity and predicted structures implied that GTAP-1 and −2 may have diverged from GCP proteins during evolution.

### GTAP-1 and −2 colocalize with γ-tubulin in the PCM throughout the cell cycle

The subcellular localization of GTAP-1 and −2 was examined using embryos expressing GFP::GTAP-1 or GFP::GTAP-2 along with mCherry::γ-tubulin and mCherry::histone. In early embryos, GFP::GTAP-1 and GFP::GTAP-2 colocalized with mCherry::γ-tubulin in the PCM throughout the cell cycle (Fig. 2 A,B). The foci areas and relative fluorescence intensities of GFP::GTAP-1 and GFP::GTAP-2 signals at centrosomes increased synchronously with γ-tubulin during centrosome maturation. At the end of telophase, the majority of GFP::GTAP-1 and GFP::GTAP-2 had dispersed to the cytoplasm, but small amounts remained at centrosomes as small foci during interphase, coinciding with γ-tubulin (Fig. 2A,B). Thus, GTAP-1 and −2 colocalized with γ-tubulin throughout the cell cycle.

**Fig. 2.**
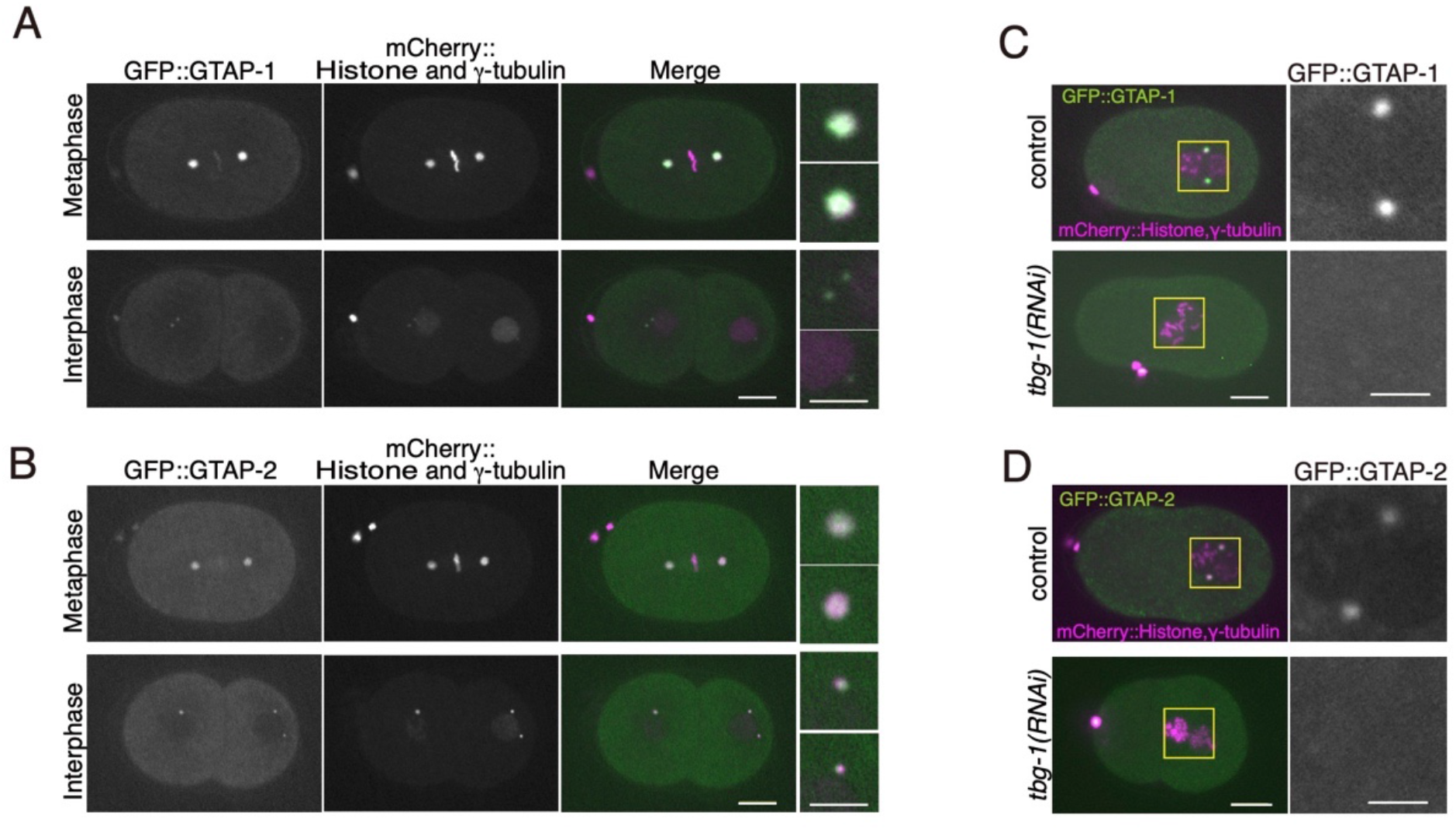
GTAP-1 and GTAP-2 localize at centrosomes in a γ-tubulin-dependent manner. (A– D) Confocal micrographs of GFP::GTAP-1 (A and C) and GFP::GTAP-2 (B and D) together with mCherry::histone H2B and mCherry::TBG-1/γ-tubulin. GTAP-1 (A) and GTAP-2 (B) colocalized with γ-tubulin in one-cell embryos throughout the cell cycle. GTAP-1 (C) and GTP-2 (D) were undetectable at centrosomes in *tbg-1/γ-tubulin (RNAi)* one-cell embryos. Scale bars indicate 10 μm in whole-embryo images and 2.5 μm in magnified images.

In *tbg-1/γ-tubulin(RNAi)* embryos, GFP::GTAP-1 (5/6 embryos) and GFP::GTAP-2 (17/18 embryos) signals were undetectable at centrosomes (Fig. 2 C, D). Therefore, the centrosomal localization of GTAP-1 and −2 was dependent on γ-tubulin, similar to the γTuSC components GIP-1/GCP3 and GIP-2/GCP2.

### GTAP-1 and −2 contribute to the efficient recruitment of γ-tubulin to centrosomes

The loss-of-function phenotypes of *gtap-1* and/or *−2* in early embryos were examined by RNAi-mediated knockdown. The *gtap-1(RNAi)*, *gtap-2(RNAi)*, and *gtap-1(RNAi);gtap-2(RNAi)* embryos were viable, but the amount of centrosomal γ-tubulin was reduced (Fig. 3 A,B). The fluorescence intensity of centrosomal GFP::γ-tubulin at metaphase in embryos at the one-cell stage was reduced to 51% (P<0.0001, n=12) in *gtap-1(RNAi)* embryos, 32% (P<0.0001, n=11) in *gtap-2(RNAi)* embryos, and 64% in *gtap-1(RNAi);gtap-2(RNAi)* embryos (P<0.001, n=10), compared with the control (n=33) (Fig. 3B). Western blotting revealed that the level of γ-tubulin was not reduced in these embryos (Fig. S2), indicating that GTAP-1 and −2 did not affect the cellular level or stability of γ-tubulin. The fact that the double depletion did not further reduce the amount of γ-tubulin at centrosomes suggested that GTAP-1 and −2 likely function in the same pathway for the efficient recruitment of γ-tubulin to centrosomes in embryos.

**Fig. 3.**
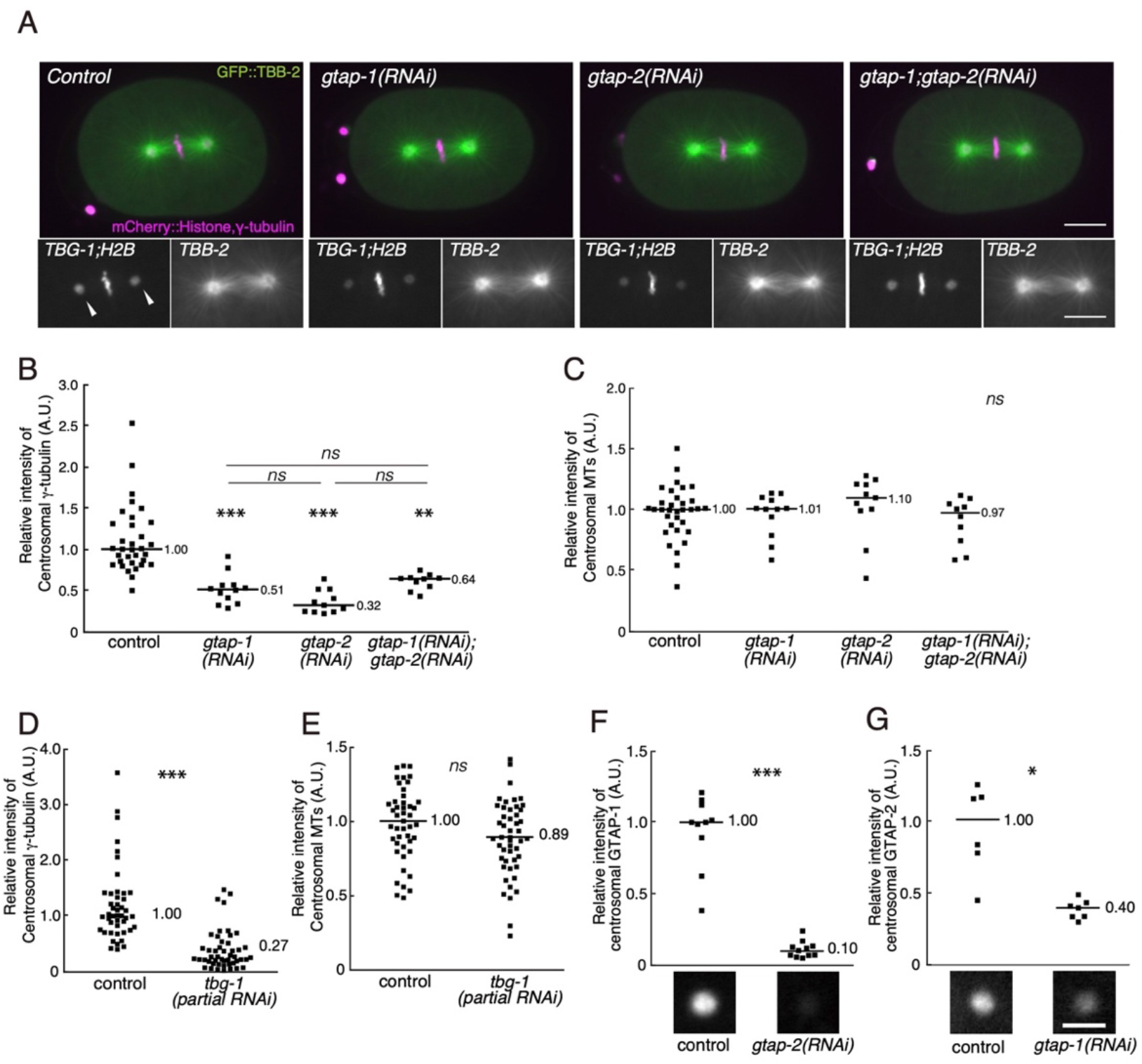
GTAP-1 and GTAP-2 contribute to the centrosomal localization of γ-tubulin. (A) Confocal micrographs of GFP::TBB-2/β–tubulin, mCherry::TBG-1/γ-tubulin, and mCherry::histone H2B in the control, *gtap-1(RNAi)*, *gtap-2(RNAi)*, and *gtap-1;gtap-2(RNAi)* embryos. Arrowheads in single-channel images indicate centrosomal signals of γ-tubulin on both sides of chromosomes. Scale bars, 10 μm. (B–G) Quantification of centrosomal signals of the GFP- or mCherry-tagged proteins at the one-cell metaphase stage. Centrosomal signals are shown for mCherry::γ-tubulin (B) and GFP::β–tubulin (C) in control, *gtap-1(RNAi)*, *gtap-2(RNAi)*, and *gtap-1;gtap-2(RNAi)* embryos. Centrosomal signals are shown for mCherry::γ-tubulin (D) and GFP::β–tubulin (E) in control and *tbg-1(partial RNAi)* embryos. (F and G) Interdependency of centrosomal localization of GTAP-1 and GTAP-2. (F) GFP::GTAP-1 in the control and *gtap-2(RNAi)* embryos. (G) GFP::GTAP-2 in the control and *gtap-1(RNAi)* embryos. Bars indicate the average fluorescence intensity for the indicated genotypes. Statistical analysis was carried out with the U-test for two groups and Tukey-Kramer multiple comparison test for four groups; *0.001<*p*<0.01, **0.0001<*p*<0.001, \*\*\**p*<0.0001. Scale bars, 2.5 μm.

### Reduction of GTAP-1 and −2 does not affect the amount of centrosomal MTs

Although centrosomal γ-tubulin was significantly reduced, the amount of centrosomal MTs was unaffected in one-cell embryos of *gtap-1(RNAi)* (101% of mean of control, n=12, P>0.1), *gtap-2(RNAi)* (110% of mean of control, n=11, P>0.1), or *gtap-1(RNAi);gtap-2(RNAi)* (97% of mean of control, n=10, P>0.1) embryos compared with the control (n=33) (Fig. 3A,C). This discrepancy between the amount of γ-tubulin and MTs at centrosomes could be explained if a large portion of centrosomal γ-tubulin is inactive for MT nucleation. To test this hypothesis, γ-tubulin was partially depleted by RNAi [*tbg-1(partial RNAi)*], and γ-tubulin and MTs surrounding centrosomes were quantified using mCherry::γ-tubulin and GFP::β-tubulin, respectively (Fig. 3 D,E). In *tbg-1(partial RNAi)* embryos, even when centrosomal γ-tubulin was reduced to ~30%, centrosomal MT content was not substantially lower (89% of mean of control, n=48, P>0.05) than that in control embryos (n=45). Moreover, bipolar spindle formation and chromosome segregation were not affected in these embryos (100%, n=48). Thus, centrosomes in one-cell embryos contained excessive γ-tubulin, and the reduction in γ-tubulin upon the loss of GTAP-1 and −2 was sufficient for MT nucleation to assemble functional mitotic spindles.

### Mutual dependency of centrosomal localization between GTAP-1 and GTAP-2

In *gtap-2(RNAi)* one-cell embryos, the level of centrosomal GFP::GTAP-1 at metaphase was reduced to 10% (P<0.01, n=11) of that in control embryos (n=10) (Fig. 3 F). Similarly, in *gtap-1(RNAi)* one-cell embryos, centrosomal GFP::GTAP-2 at metaphase was reduced to 40% (P<0.01, n=7) of that in control embryos (n=7) (Fig. 3G). Western blotting revealed that the amount of GTAP-1 and GTAP-2 was not reduced in *gtap-2(RNAi)* and *gtap-1(RNAi)* embryos, respectively (Fig. S2). Thus, centrosomal localization of GTAP-1 and −2 was partially dependent on each other, although GTAP-1 was more dependent on GTAP-2.

### GTAP-1 and −2 affect PCM integrity

Although the amount of centrosomal MTs was barely affected in *gtap-1(RNAi)*, *gtap-2(RNAi)*, and *gtap-1(RNAi);gtap-2(RNAi)* embryos, their interaction with the PCM seemed to be altered. In wild-type embryos at the end of telophase, the dynein-dependent cortical MT pulling force mediates fragmentation of spindle poles (Grill et al., 2001; Grill et al., 2003; Labbe et al., 2004; Pecreaux et al., 2006; Kozlowski et al., 2007), and γ-tubulin in the PCM scatters into the cytoplasm. In *gtap-1(RNAi)*, *gtap-2(RNAi)*, and *gtap-1(RNAi);gtap-2(RNAi)* embryos, fragmentation of the spindle pole occurred earlier and more drastically than expected (Fig. 4A and S3). Concurrently, PCM, as visualized by GFP::SPD-5, was also precociously fragmented (Fig. 4B). In *tbg-1(partial RNAi)* embryos, this fragmentation of the spindle pole was not observed; instead, the signal of the spindle pole gradually decreased (Fig. 4A and S3). These observations indicated that exaggerated fragmentation of the spindle pole in GTAP-1-and/or GTAP-2-depleted embryos was not simply caused by a reduction in the level of centrosomal γ-tubulin; rather, depletion of GTAP-1 and −2 seemed to affect PCM integrity, by, for example, affecting the interaction between γTuC and PCM scaffold components. An alternate possibility is that the dynamics of MTs to generate the cortical pulling force might differ between the MTs nucleated from intact γTuC and those nucleated from an incomplete γTuC without GTAP-1 and 2.

**Fig. 4.**
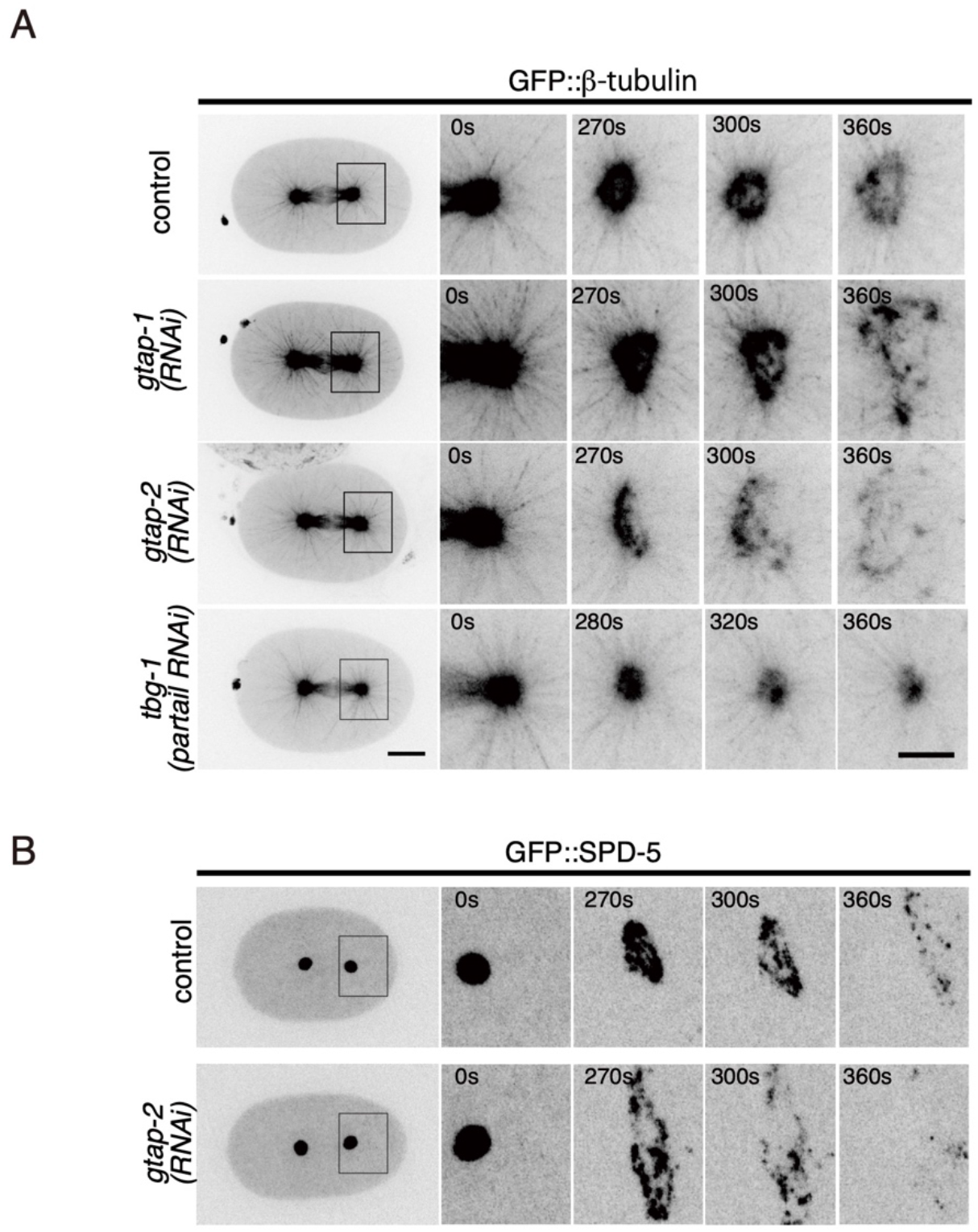
GTAP-1 and GTAP-2 affect centrosome disassembly in late mitosis. (A) Time-lapse confocal micrographs of GFP::β-tubulin in the control, *gtap-1(RNAi)*, *gtap-2(RNAi)*, and *tbg-1*/ γ *-tubulin (partial RNAi)* embryos. (B) Time-lapse confocal micrographs of GFP::SPD-5 in the control and *gtap-2(RNAi)* embryos. The left-most panels show images of whole embryos (scale bars, 10 μm). The right panels are magnified views of the centrosomal regions in the time-lapse images (scale bars, 5 μm). More than six embryos were observed for each genotype, and representative images were shown. To avoid the effect of photobleaching in different embryos, all embryos were monitored in brightfield mode until metaphase when the fluorescence imaging was initiated. Times are relative to metaphase.

### GTAP-1 and GTAP-2 contribute differently to the development of the hermaphrodite germline

To examine the developmental phenotypes, null mutants of *gtap-1* and *gtap-2* were constructed by CRISPR/Cas9. In each of the null mutants, namely *gtap-1(tj84)* and *gtap-2(tj92)*, centrosomal γ-tubulin was reduced by ~50% in one-cell embryos, as was the case in the *gtap-1(RNAi)* and *gtap-2(RNAi)* embryos (Fig. S4), and could be maintained as homozygotes, suggesting that GTAP-1 and −2 are not essential for development and fertility. However, the fecundity of *gtap-1(tj84)* worms was severely reduced, especially in Day 2 and Day 3 adults (Fig. 5A). Correspondingly, the embryonic lethality of the *gtap-1* mutant increased with age, i.e., up to ~50% by Day 3. The variability of the size and shape of *gtap-1(tj84)* embryos also increased with age (Fig. 5C). In contrast, mutant *gtap-2(tj92)* did not show embryonic lethality (Fig. 5B), and the proportion of embryos having abnormal size and shape was lower than that for *gtap-1(tj84)* (Fig. 5C).

**Fig. 5.**
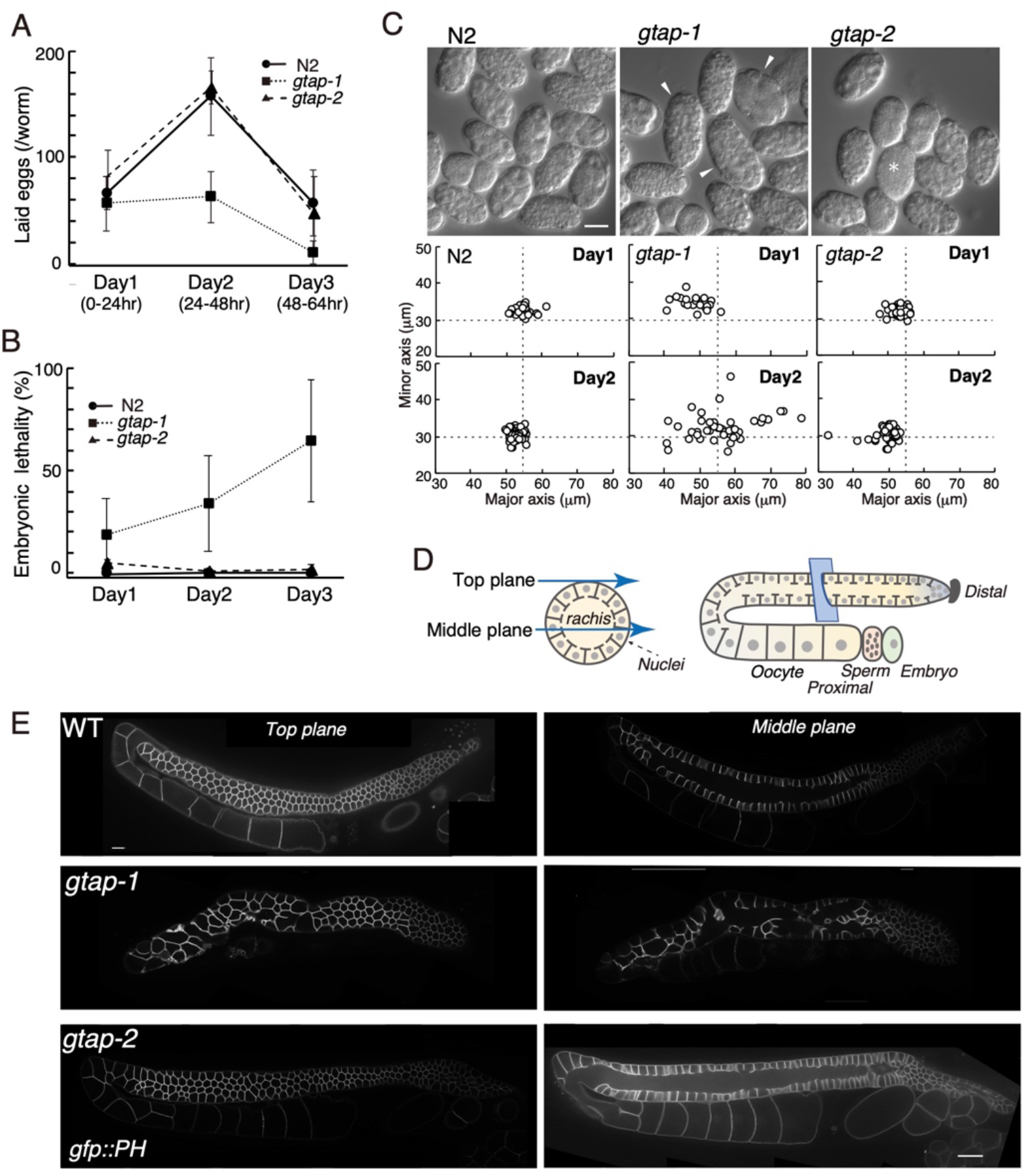
GTAP-1 is required for organization of the germ-cell compartment. (A) Number of laid eggs in 24-h windows in the control (N2), mutant *gtap-1*, and mutant *gtap-2*. (B) Embryonic lethality of the eggs in (A). Embryonic lethality of the *gtap-1* mutant increased in an age-dependent manner. (C) Egg size variability of the *gtap-1* mutant. Differential interference contrast micrographs of eggs laid by Day 2 adult worms (upper images) and the sizes of the eggs laid by Day 1 and Day 2 adults (lower graphs). White arrowheads indicate abnormally sized embryos, and the asterisk indicates an unfertilized egg. (D) Schematic drawing of a *C. elegans* adult hermaphrodite gonad (modified from Zhou et al., 2009). (E) Confocal micrographs of adult hermaphrodite gonads that expressed GFP::PH (plasma membrane marker). The germline in the *gtap-1* mutant was morphologically abnormal. Top (left) and middle (right) focal planes of the gonad are shown. Scale bars in panels C and E, 20 μm.

The morphologically abnormal eggs and reduction of brood size in the *gtap-1(tj84)* mutant implied defects in oogenesis. In adult hermaphrodites, the distal gonad is syncytial, and each meiotic nucleus is compartmentalized with the membranes organized in a honeycomb-like structure. Each compartment is partially open and connected to the large cytosolic region called the rachis (Fig. 5D). The organization of the adult hermaphrodite germline was analyzed using a strain in which the germ-cell membrane was labeled by GFP::PH (Pleckstrin-Homology) domain (Fig. 5E). In the gonad of *gtap-1(tj84)* adults, the honeycomb-like structure of the germ-cell membrane was disorganized, and the size of the compartment was highly variable. On the other hand, the honeycomb-like structure of the *gtap-2(tj92)* adult germline was only slightly perturbed. The *gtap-1(tj84);gtap-2(tj92)* double mutant had more severe and a wider range of phenotypes than either single mutant (a small brood size of only 6.3 eggs/worm, 49.3% embryonic lethality, N=11; low frequency of Dumpy phenotype or rupture of adult worms). These results indicated that, whereas GTAP-1 plays a more important role in the adult germline, GTAP-1 and −2 function redundantly in various tissues.

### Both GTAP-1 and GTAP-2 colocalize at the plasma membrane in the germline where they contribute to the efficient recruitment of γ-tubulin

Next, the subcellular localization of GTAP-1 and GTAP-2 in the hermaphrodite germline was analyzed using strains in which each endogenous GTAP was tagged with GFP by CRISPR/Cas9 (Fig. 6A). Both GFP-labeled proteins localized at centrosomes and at the plasma membrane of the adult germline, and GFP::GTAP-2 was also detected in the cytoplasm. These localization patterns at centrosomes and plasma membrane in the adult germline were similar to that of γ-tubulin. The γ-tubulin at the plasma membrane is required for the nucleation of MTs, involving the positioning of germ-cell nuclei in each compartment of the syncytial gonad (Zhou et al., 2009). Therefore, we next examined whether plasma-membrane localization of γ-tubulin was affected in the *gatp-*1*(tj84)* and *gatp-2(tj92)* mutants using a strain co-expressing mCherry::γ-tubulin and mCherry::histone H2B. The amount of γ-tubulin at the plasma membrane was quantified based on the relative intensity of mCherry::γ-tubulin using mCherry::histone H2B as the internal control (Fig. 6B). The average relative intensity (γ-tubulin::histone) was 0.37±0.10 in control worms (Fig. 6B). In the *gtap-1(tj84)* and *gtap*-2*(tj92)* mutants, however, the relative intensity of γ-tubulin was significantly reduced to 0.18±0.069 and 0.22±0.086, respectively (Fig. 5B), suggesting that GTAP-1 and −2 contribute to the efficient recruitment of γ-tubulin to the plasma membrane in the adult germline.

**Fig. 6.**
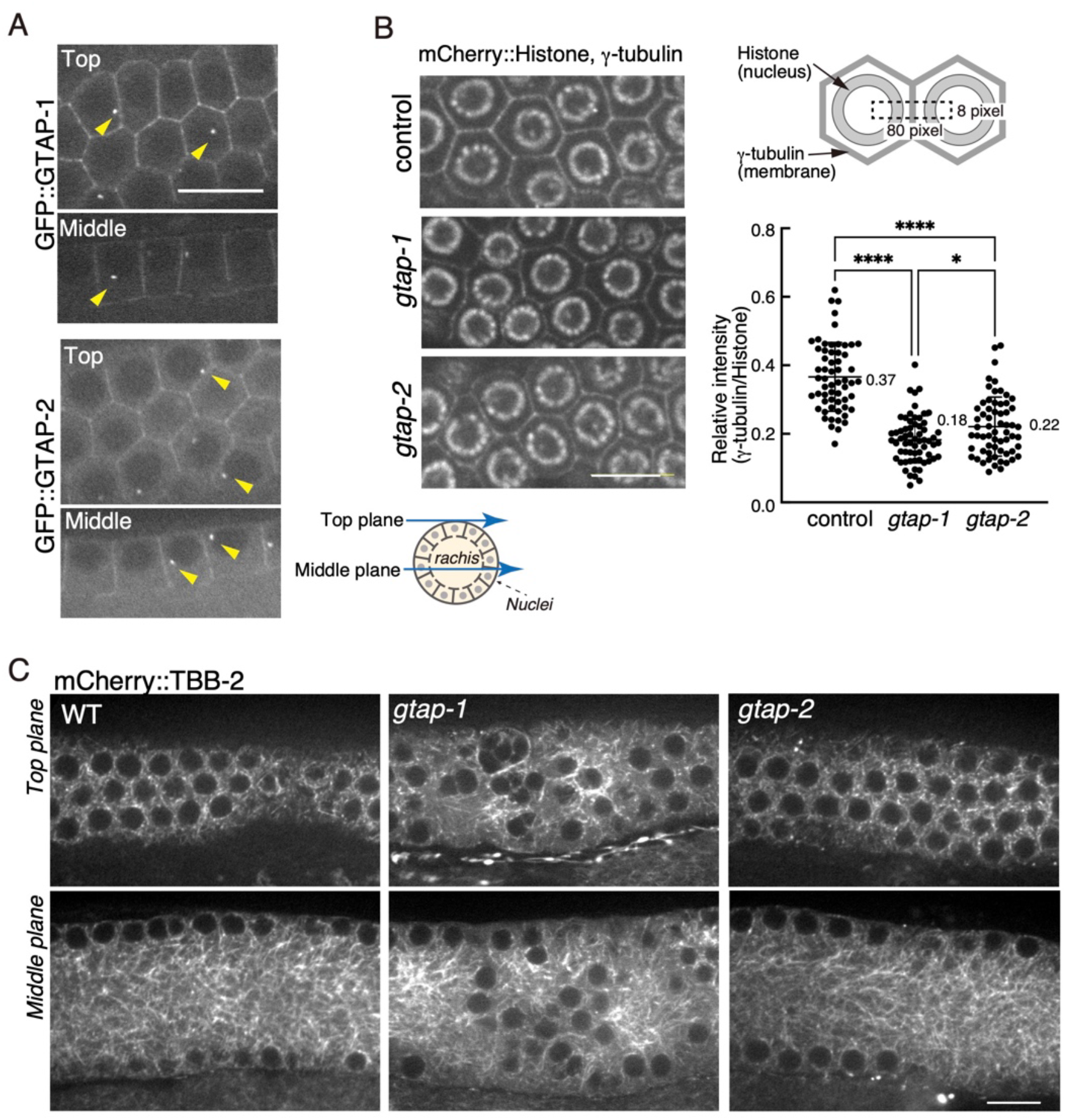
GTAP-1 is required for the positioning of germ-cell nuclei in the adult hermaphrodite gonad. (A) Endogenous GFP::GTAP-1 and GFP::GTAP-2 localized to the germ-cell membrane and centrosomes (yellow arrowheads). (B) The localization of mCherry::γ-tubulin on the germline membrane was reduced in *gtap-1* and *gtap-2* mutants. The fluorescence intensity of mCherry::γ-tubulin on the plasma membrane was calculated as the intensity relative to the average intensity of mCherry::histone in the dotted square indicated in the schematic diagram. Statistical analysis was carried out with the Tukey-Kramer multiple comparison test for three groups; *0.001<*p*<0.01, **0.0001<*p*<0.001, \*\*\**p*<0.0001. (C) Confocal micrographs of adult hermaphrodite gonads that expressed mCherry::TBB-2. The MT array around the nuclei was disrupted, and some of the nuclei were mislocated in the rachis in the *gtap-1* mutant. In the *gtap-2* mutant, the MT array surrounding the nuclei was not substantially affected. Scale bars, 10 μm.

### GTAP-1 is required for the organization of the MT array around germline nuclei

Because the amount of γ-tubulin at the plasma membrane in the germline was decreased in each of the *gtap-1(tj84)* and *gtap*-2*(tj92)* mutants, MTs in the germline were observed by live imaging using mCherry::tbb-2. In wild-type and *gtap-2(tj92)* adult hermaphrodites, MTs were highly organized around germ-cell nuclei located at the periphery of the syncytial gonad (Fig. 6C). In the *gatp-1(tj84)* mutant, however, the distribution of MTs surrounding nuclei was uneven, the location of nuclei was perturbed, and some nuclei were detached from the peripheral compartments and detected in the rachis (Fig. 6C). Thus, even though both GTAP-1 and −2 contributed to the recruitment of γ-tubulin to the plasma membrane in the germline, GTAP-1 played a much larger role than GTAP-2 in the organization of MT arrays.

### GTAP-1 and −2 are associated with γTuC *in vivo*

Because the embryonic and germline subcellular localizations and phenotypes strongly indicated a close link between each of GTAP-1 and −2 and γ-tubulin, the interaction of GTAP-1 and −2 with γTuSC components was examined by immunoprecipitation (Fig. 7A). The immunoprecipitation was performed with anti-GFP using extracts of embryos of the control strain N2 and of strains expressing either GFP::γ-tubulin, GFP::GTAP-2, or GFP::GTAP-1. As expected, from the extract of the strain that expressed GFP::γ-tubulin, GTAP-1 and −2 co-immunoprecipitated along with GIP-1/GCP3 and GIP-2/GCP2. Conversely, from the extracts of the strains that expressed GFP::GTAP-1 or GFP::GTAP-2, all components of the γTuSC (γ-tubulin, GIP-1/GCP3 and GIP-2/GCP2) as well as GTAP-2 or −1 co-precipitated, respectively. These results indicated that GTAP-1 and −2 interact with the γTuC *in vivo*.

**Fig. 7.**
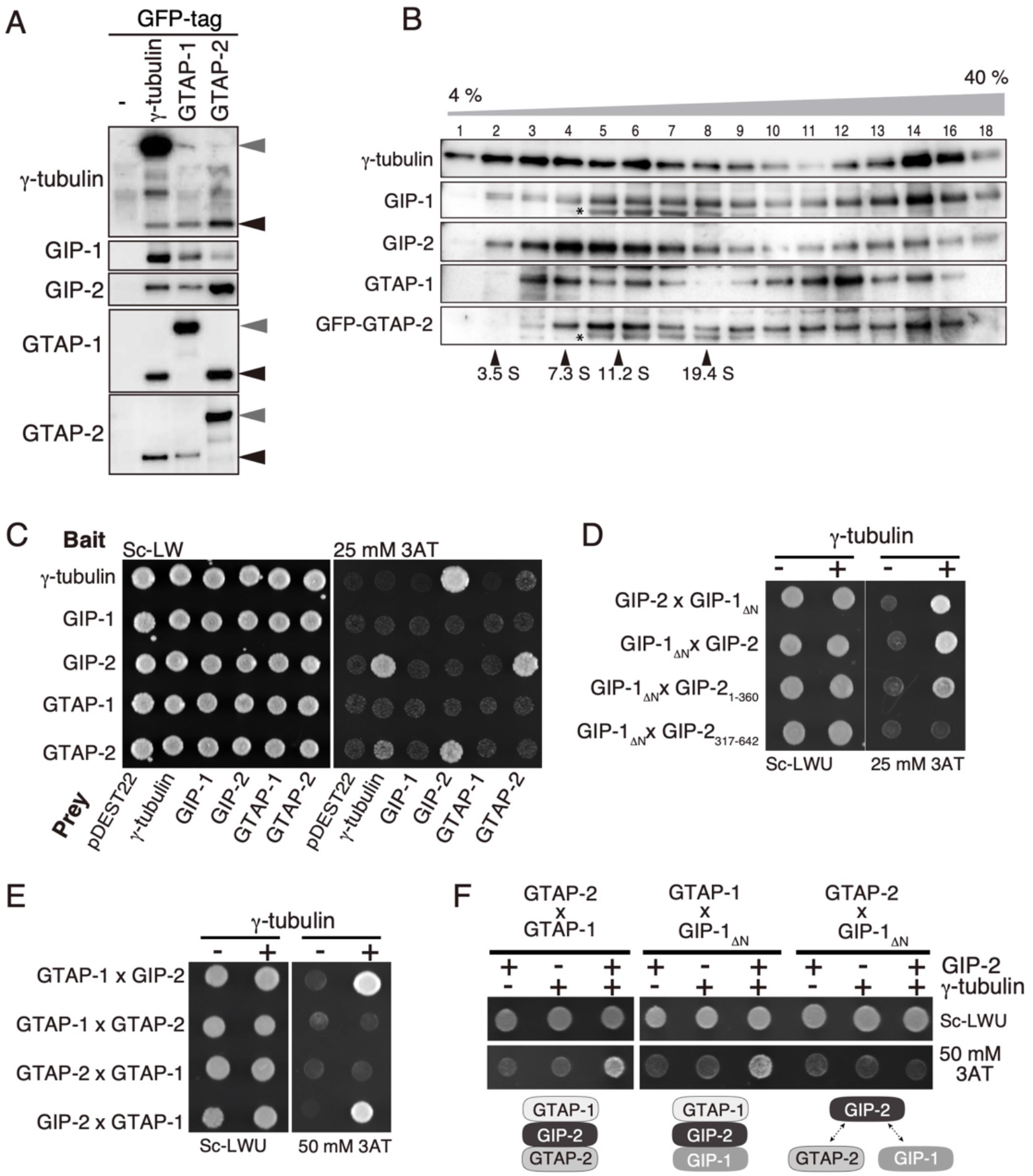
Association of GTAP-1 and GTAP-2 with the components of γTuC. (A) Co-immunoprecipitation of GTAP-1 and −2 with the γTuC components. Components of the γTuC were co-immunoprecipitated from extracts of embryos expressing GFP-tagged γ-tubulin (SA303), GTAP-1 (SA536), or GTAP-2 (SA526) using anti-GFP. Immunoprecipitates were analyzed by western blotting with an antibody specific for each protein. Gray arrowheads indicate the GFP-tagged exogenous proteins, and black arrowheads indicate endogenous proteins. (B) Sucrose gradient ultracentrifugation for 4 h. Embryonic extracts prepared from strain SA526 (GFP::GTAP-2) were fractionated on a 4–40% sucrose gradient. The indicated fractions were analyzed by western blotting with an antibody specific for each protein or against GFP to detect GFP::GTAP-2. The following molecular standards were used: ovalbumin (3.5 S), aldolase (7.3 S), catalase (11.2 S), and thyroglobulin (19.4 S). Asterisks indicate non-specific bands. (C–F) Yeast two-hybrid assays with GTAP-1, GTAP-2 and γTuC components. (C) Detection of interactions among GTAP-2, GIP-2/GCP2, and γ-tubulin. (D) Detection of the interaction between GIP-1_ΔN_ (N-terminally truncated residues 293–891) with GIP-2 in the presence of γ-tubulin. (E) Detection of the interaction between GTAP-1 and GIP-2 in the presence of γ-tubulin. Interaction of GTAP-2 with GIP-2 was not detected. (F) Detection of the interaction of GTAP-1 with GIP-1_ΔN_, or GTAP-2 in the presence of both γ-tubulin and GIP-2. Interaction between GIP-1_ΔN_and GTAP-2 was not detected.

Endogenous GTAP-1 was not detected among the proteins that co-immunoprecipitated with GFP::GTAP-1; similarly, endogenous GTAP-2 was not detected among proteins that co-immunoprecipitated with GFP::GTAP-2. Thus, we speculated that γTuC associates with one molecule of GTAP-1 and one molecule of GTAP-2.

To further examine whether γTuCs in *C. elegans* contain GTAP-1 and/or GTAP-2, embryo extracts were subjected to sucrose gradient sedimentation. Two distinct peaks of γ-tubulin-containing fractions were detected (fractions 1–9 and 12–16) (Fig. 6B), whose sizes roughly corresponded to that of γTuSC and γTuRC in other organisms (estimated S-values were 4–15 S and 25–35 S, respectively) (Stearns and Kirschner, 1994; Zheng et al., 1995; Murphy et al., 1998; Oegema et al., 1999; Anders et al., 2006). However, the separation of the two peaks was less clear than what has been observed in experiments with other organisms, and both γ-tubulin peaks contained GIP-1/GCP3, GIP-2/GCP2, GTAP-1, and GTAP-2. We speculated that γTuCs in *C. elegans* are heterogeneous, including various numbers of γTuSCs with GTAP-1 and/or GTAP-2.

### GTAP-1 and −2 directly interact with components of the γTuSC

Because GTAP-1 and −2 were found to associate with γTuC *in vivo*, the direct interaction of GTAP-1 and −2 with γTuSC components was examined using yeast two-hybrid assays (Fig. 7C and Table S1). GTAP-2 interacted strongly with GIP-2/GCP2 and weakly with γ-tubulin (Fig. 7C). GIP-2/GCP2 interacted strongly with γ-tubulin as well as GTAP-2 (Fig. 7C). Interactions between GTAP-1 and other components were not detected (Fig. 7C).

Because the predicted interactions of GIP-1/GCP3 with other γTuSC components (γ-tubulin and GIP-2/GCP2) were not detected in this assay, we speculated that some interactions might require more than two γTuC components. Therefore, interactions between three components were examined using a modified yeast two-hybrid assay. GCP proteins, including GIP-1/GCP3 and GIP-2/GCP2, have the grip1 and grip2 motifs that interact with adjacent GCP proteins and γ-tubulin, respectively. In subsequent assays, the core region of GIP-1/GCP3 (271– 891, termed GIP-1_ΔN_) containing the grip1 and grip2 motifs was used. When γ-tubulin was co-expressed, GIP-1_ΔN_ interacted with GIP-2 (Fig. 7D). GIP-2/GCP-2 (1–360) lacking the grip2 motif still interacted with GIP-1_ΔN_ in a γ-tubulin-dependent manner, whereas GIP-2/GCP-2 (317– 642) lacking the grip1 motif did not. Thus, this ternary complex is likely to require interactions between γ-tubulin and GIP-1_ΔN_ and between GIP-1 _ΔN_ and the grip1 motif of GIP-2/GCP2 (Fig. 7 D).

In a modified yeast two-hybrid assay with γ-tubulin, GTAP-1 interacted with GIP-2/GCP-2 and GIP-2/GCP-2 (1–360), whereas interaction between GTAP-1 and GTAP-2 was not detected (Fig. 7E and Table S1). This result indicated that the GTAP-1-GIP-2/GCP2-γ-tubulin ternary complex requires interactions between γ-tubulin and GTAP-1 and between GTAP-1 and the grip1 motif of GIP-2/GCP2. These interactions were similar to those observed for the GIP-1/GCP3-GIP-2/GCP2-γ-tubulin ternary complex.

Because GTAP-1, GTAP-2, and GIP-1/GCP3 were found to interact with GIP-2/GCP2, we next investigated whether these three proteins compete for binding to the same site in GIP-2/GCP2 using a modified yeast two-hybrid assay in which γ-tubulin and GIP-2/GCP2 were co-expressed in addition to the bait and prey proteins. In the presence of γ-tubulin and GIP-2/GCP2, the interactions (either direct or indirect) between GTAP-1 and −2 or between GTAP-1 and GIP-1/GCP3 were detected, but the interaction between GTAP-2 and GIP-1/GCP3 was not (Fig. 7F). These results indicated that GTAP-2 competes with GIP-1/GCP3 for binding to GIP-2/GCP2, whereas GTAP-1 does not compete with either GIP-1/GCP3 or GTAP-2.

### MZT-1/MOZART1 plays a more crucial role than GTAP-1 and −2 in the centrosomal recruitment of the γTuC in early embryos

MOZART1 is a broadly conserved small protein involved in the γTuC and is essential for spindle assembly in many organisms. *C. elegans* MOZART1, MZT-1, accumulates in the PCM and is required for the interaction between the N-terminus of SPD-5 and γ-TuSC (GIP-1/GCP3, GIP-2/GCP2, and γ-tubulin) (Sallee et al., 2018; Ohta, et al., 2021). To compare the roles of GTAP-1 and −2 with that of MZT-1, we examined its localization and loss-of-function phenotypes in early embryos by RNAi-mediated knockdown. As previously reported, MZT-1 accumulated in the PCM, and its recruitment was dependent on γ-tubulin and GIP-1/GCP3 (Fig. 8A). Whereas the depletion of GTAP-1 and/or −2 did not cause a significant defect in chromosome segregation (Fig. 3), depletion of MZT-1 resulted in severe defects of both spindle formation and chromosome segregation (Fig. 8B). The PCM localization of γ-tubulin and GIP-1/GCP3, but not of SPD-5, was significantly reduced in MZT-1-depleted embryos (Fig. C). The amount of centrosomal γ-tubulin was reduced to ~6% of the control (p<0.0001, n=15), resulting in a significant reduction of centrosomal MTs (72% of the control, p<0.001, n=15) (Fig. 8D,E). Thus, MZT-1 in early embryos plays a more crucial role than GTAP-1 or −2 in the centrosomal localization of γTuC.

**Fig. 8.**
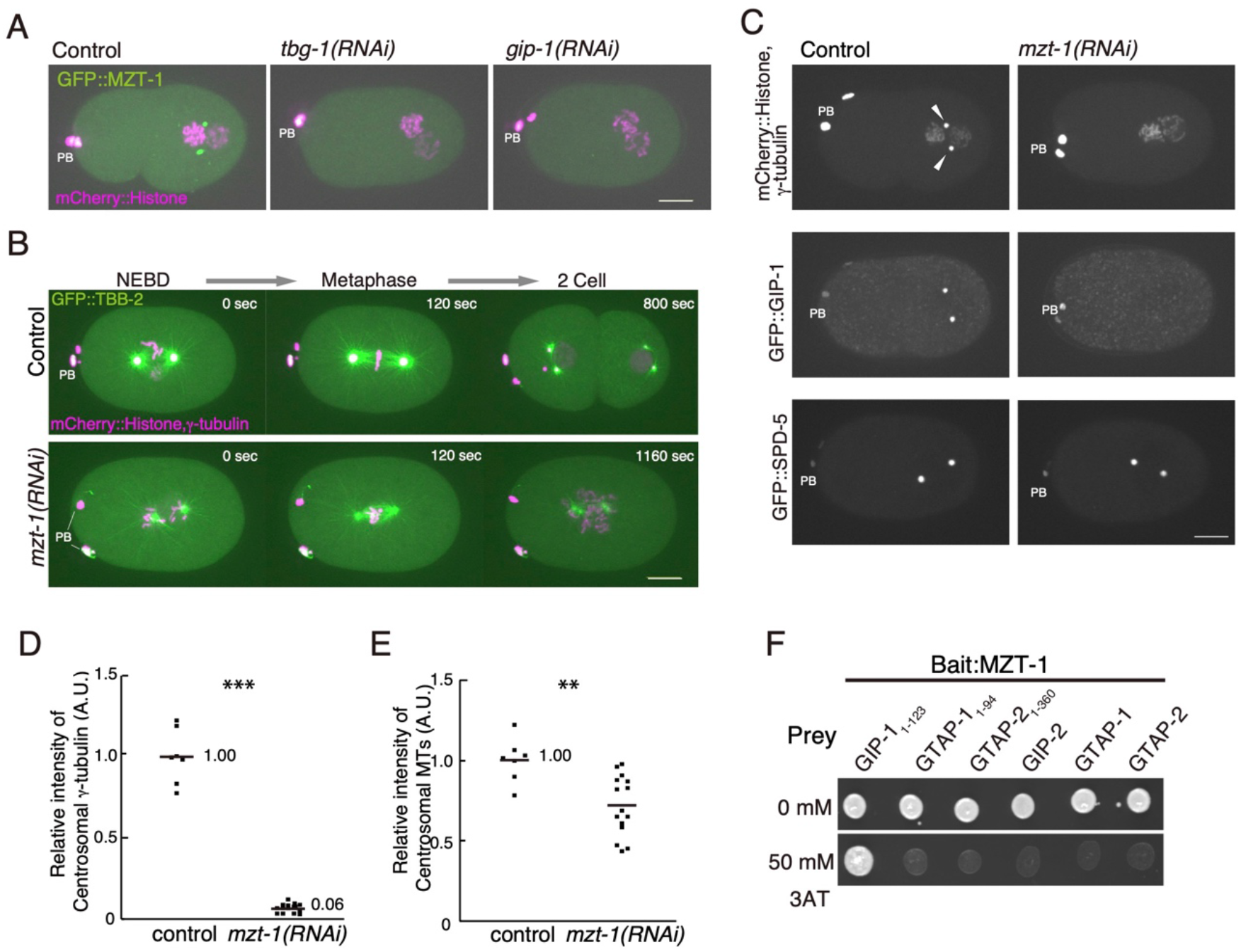
MZT-1 is essential for centrosomal recruitment of γ-tubulin in early embryos. (A) γTuC-dependent centrosomal localization of GFP::MZT-1. Confocal micrographs show GFP::MZT-1 and mCherry::histone H2B at the pronuclei meeting stage. Scale bars, 10 μm. (B) Severe mitotic spindle defects in *mzt-1(RNAi)* embryos. Time-lapse confocal micrographs are shown of GFP::TBB-2/β-tubulin, mCherry::TBG-1/γ-tubulin and mCherry::histone H2B from nuclear envelope breakdown (NEBD) to 2-cell stages. PB indicates the polar body. (C) MZT-1-dependent centrosomal localization of γTuC. Confocal micrographs are shown of mCherry::TBG-1/γ-tubulin and mCherry::histone H2B, GFP::GIP-1 and GFP-SPD-5 at the pronuclei meeting stage in the control and *mzt-1(RNAi)* embryos. White arrowheads indicate the centrosomal signals of mCherry::TBG-1/γ-tubulin. (D and E) Quantification of centrosomal signals of the fluorescent proteins at the one-cell metaphase stage. (D) mCherry::TBG-1/γ-tubulin. (E) GFP::TBB-2/β-tubulin. Statistical analysis was carried out with the U-test; *0.001<*p*<0.01, **0.0001<*p*<0.001, \*\*\**p*<0.0001. (F) Detection of the interaction between MZT-1 and GIP-1/GCP3 (N-terminus) by a yeast two-hybrid assay. Interaction of MZT-1 with GIP-2/GCP2, GTAP-1 (full-length and N-terminus), and GTAP-2 (full-length and N-terminus) was not detected.

In *Arabidopsis* and other organisms, MOZART1 orthologs bind the N-terminal region of GIP-1/GCP3, except for the human MOZART1 ortholog, which likely binds the N-terminal region of GCP2, GCP3, GCP5, and GCP6 (Janski et al., 2012; Nakamura et al., 2012; Dhani et al., 2013; Cota et al., 2017). A yeast two-hybrid assay revealed that *C. elegans* MZT-1 bound to the N-terminal region of GIP-1/GCP3 but not GIP-2/GCP2, GTAP-1 or GTAP-2 (Fig. 8F). Because MZT-1 was required for the centrosomal localization of GIP-1/GCP3 (Fig. 8C), this interaction might be essential for γTuSC localization to centrosomes.

### Phylogenetic analysis of γTuC components

As described above, GTAP-2 contains a region that has only modest sequence similarity to part of the grip-2 motif conserved in GCP2-6 proteins in other organisms. The predicted protein structure of GTAP-1 is similar to the common GCP structure, even though the sequence similarity was undetectable (Fig. 1C). These findings implied the possibility that GTAP-1 and −2 might be highly divergent versions of GCP4–6. To understand how the composition of γTuC has evolved, the phylogeny of each γTuC component was analyzed for 28 species (12 metazoa including 7 nematode species, 3 protozoa, 9 fungi, and 4 plants) (Fig. 9A), which led to the following findings.

**Fig. 9.**
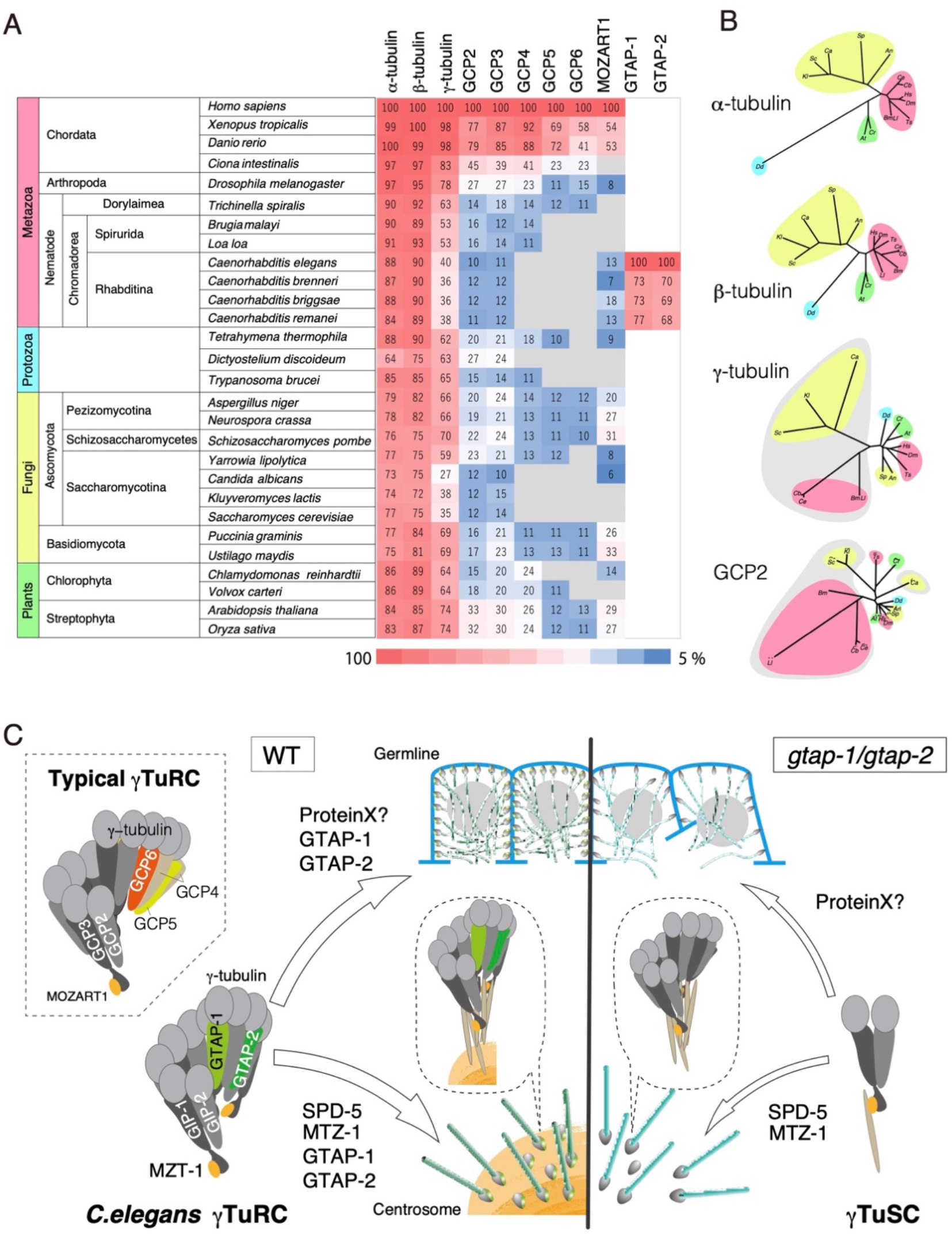
Phylogenetic analysis of γTuC components. (A) Phylogenetic analysis of the tubulins and components of γTuC among eukaryotes. The scores indicate the amino-acid sequence identity between the human protein and its orthologs. (B) Phylogenetic trees for tubulins and GCP2. Colors correspond to Metazoa (red), Protozoa (blue), Fungi (yellow) and Plants (green), respectively. Each gray area in the γ-tubulin and GCP2 drawings indicates that the organisms lacked GCP4–6. (C) Hypothetical models for the roles of GTAP-1 and GTAP-2 in γTuRC functions in *C. elegans* based on this study. The γTuRC with GTAP-1 and GTAP-2 is efficiently recruited to centrosomes and the plasma membrane in the germline. In the absence of GTAP-1 and GTAP-2, a reduced amount of γTuC is still recruited to centrosomes and the plasma membrane in the germline. GTAP-1 plays a more pivotal role than GTAP-2 in MT organization in the adult germline.

First, all eukaryotes that were examined contained γTuSC components (γ-tubulin, GCP2 and GCP3), and at least some species in all four groups (metazoa, protozoa, fungi and plants) contained GCP4–6, suggesting that the canonical γTuRC composition was established at the emergence of eukaryotes, i.e., before diverging into these four groups. Second, at least twice during evolution, some of the γTuRC-specific components were lost from the genome. *Saccharomyces cerevisiae* and two other fungi that belong to *Saccharomycotina* (*Candida albicans* and *Kluyveromyces lactis*) do not have GCP4–6, whereas the majority of other fungi have the complete set. Also, within nematode species, whereas *Trichinella spiralis* has the complete set of GCP4–6, species in Chromadoria (*Brugia malayi*, *Loa loa* and four *Caenorhabditis* species including *C. elegans*) lack some of them. GTAP-1 and −2 are highly conserved in *Caenorhabditis* species although their phylogenetic relationship with GCP4–6 is unclear. Third, in the species that had lost some of GCP4–6, the evolution of the remaining components was accelerated (Fig. 9B). Whereas the phylogenetic trees for α- and β-tubulins are consistent with the phylogeny of the organisms, in the phylogenetic tree for γTuSC components, all organisms that lack some of GCP4–6 are separated from those with the complete sets. These data imply that structural constraints of the γTuSC components are released by the loss of GCP4–6 to adopt a new stable state of γTuRC as a whole.

## DISCUSSION

In this study, we demonstrated that *C. elegans* has an unconventional composition of γTuC by identifying two γTuC components, namely GTAP-1 and GTAP-2, which are predicted to be highly divergent versions of GCP4–6 in other organisms. GTAP-1 and −2 contribute to the efficient recruitment of γ-tubulin to centrosomes in early embryos and to the plasma membrane in the germline. Unlike another γTuC component MZT-1, however, GTAP-1 and −2 are not essential for embryogenesis (Fig. 9C). GTAP-1 plays a critical role in establishing the organization of germline cells in adults, whereas GTAP-2 plays a minor role in this process (Fig. 9C). We propose that GTAP-1 and −2 contribute to the organization of centrosomal and non-centrosomal MTs in a tissue-specific manner.

### γTuC in *C. elegans*

Our results demonstrate that GTAP-1 and −2 directly interact with GIP-2/GCP2. As such, we predict that the structure of the *C. elegans* γTuC—which includes GTAP-1 and GTAP-2— will be similar to that of the γTuC region encompassing GCP4–6 in other organisms, ultimately mirroring the canonical ring-like structure of γTuC (Fig. 9C). Structural analyses of vertebrate γTuRCs have indicated that two molecules of GCP4 and one molecule each of GCP5 and GCP6 are incorporated in the half region of γTuRC (Liu et al., 2019; Wieczorek et al., 2020; Consolati, et al., 2020). The structure of the region that contains GCP4–6 is open and more flexible than the rest of the γTuRC. A phylogenic analysis implies that an ancestral nematode species of *C. elegans* lost at least one of the conventional γTuRC-specific components (GCP4–6), which led to the rapid co-evolution of the remaining γTuC components. Because the amino-acid sequence similarity between GTAP-1 and −2 with GCP4–6 was minimal, it is unclear which GCP(s) corresponds to GTAP-1 and −2.

*C. elegans* lacks several γTuRC-interacting proteins, such as NEDD-1/GCP-WD and the Augmin complex (Uehara et al., 2009), that are conserved in other organisms that have conventional γTuRC. One possibility is that these proteins might have been lost as a consequence of the alteration of the composition and γTuC structure in *C. elegans*. Further structural analysis will be needed to understand how the *C. elegans* γTuC containing GTAP-1 and −2 differs in a structural sense from the structure of conventional γTuRCs.

### GTAP-1 and −2 function during embryogenesis

Our results suggest that the role of GTAP-1 and −2 in centrosomal recruitment of γ-tubulin is distinct from that of MZT-1. MZT-1 is required for the centrosomal targeting of γTuC by mediating the interaction between γTuSC and phosphorylated SPD-5 at the mitotic PCM (Sallee et al., 2018; Ohta, et al., 2021). Although depletion of MZT-1 resulted in a ~95% reduction of centrosomal γ-tubulin, depletion of GTAP-1 and/or −2 caused a 50–70% reduction, indicating that the MZT-1 plays an essential role in centrosomal recruitment of γTuSC but GTAP-1 and −2 do not. We speculate that, although γTuSC can be recruited to the PCM via the interaction between MZT-1 and SPD-5, the γTuC containing GTAP-1 and −2 may be more stable than γTuC without them, which may contribute to the efficient accumulation of γTuC in the PCM.

Although GTAP-1 and −2 are dispensable for γTuC recruitment to the PCM, our data indicate that their absence affects the dynamics of both centrosomal MTs and the PCM at telophase. One possibility is that GTAP-1 and −2 may contribute to strengthening the connections among γTuCs and PCM proteins at centrosomes, which in turn contributes to the integrity of the PCM. Alternatively, the MTs formed from a γTuC lacking GTAP-1 and/or −2 may result in distinct dynamics at MT ends or altered interaction with MT binding proteins or motors such as dynein, which are involved in pulling forth from the cell cortex (Grill et al., 2001; Grill et al., 2003; Labbe et al., 2004; Pecreaux et al., 2006; Kozlowski et al., 2007).

### A crucial role for GTAP-1 in the germline

In the adult germline, the γTuC localizes to the gonadal membrane of syncytial gonads, and MTs emanating from the membrane are required for maintaining the germline nuclei in each honeycomb-like cell compartment (Zhou et al., 2009). GTAP-1 and −2 contribute to the efficient recruitment of the γTuC to the plasma membrane in the germline. The *gtap-1* mutant had severe morphological defects in the organization of the honeycomb-like gonadal compartment, which is similar to what has been observed for the depletion of ZYG-12, which localizes to the nuclear envelope and anchors MTs emanating from the germline membrane (Zhou et al., 2009). We speculate that the γTuC containing GTAP-1 assembles MT arrays on the plasma membrane, and anchoring these MTs to the nuclear envelope via ZYG-12 is crucial for the organization of meiotic nuclei in the compartments of the germline.

Although the amounts of γ-tubulin at the plasma membrane in the gonads was reduced to a similar extent in both the *gtap-1* and *gtap-2* mutants, only *gtap-1* had a severe phenotype. This indicates that the role of GTAP-1 is not limited to the efficient recruitment of the γTuC to the plasma membrane in the germline, and indeed it may be involved in the structural and/or functional regulation of γTuC.

The phenotype of *gtap-1* is also similar to that observed upon depletion of NOCA-1, which is another protein that localizes to the minus-end of MTs (Green et al., 2011; Wang et al., 2015). Unlike GTAP-1 and −2, however, the localization of γTuC at the germline membrane is not dependent on NOCA-1, although the localization of NOCA-1 is partially dependent on the γTuC (Wang et al., 2015). This similarity of phenotypes indicates that GTAP-1 and NOCA-1 may cooperatively regulate γTuC to assemble the MT array on the germline membrane. Further analysis will be needed to understand the interactions among γTuC, GTAP-1, GTAP-2, and NOCA-1 with regard to MT organization in the germline.

### Tissue specificity of γTuRC at non-centrosomal MTOC

Our study demonstrates that the requirement of GTAP-1 and −2 for γTuC recruitment may differ between mitotic centrosomes and the non-centrosomal MTOC (ncMTOC) at the plasma membrane in the germline. Although GTAP-1 is dispensable for embryogenesis and has nearly equivalent function to GTAP-2 with regard to centrosomal targeting of the γTuC, it plays a more crucial role than GTAP-2 in germline organization. These findings indicate that certain γTuC components play different roles between centrosomal the MTOC and ncMTOC. Similarly, human GCP6 plays a role in the localization of γTuC to keratin fibers in epithelial cells in addition to its ubiquitous role in γTuRC assembly (Oriolo et al., 2007; Liu et al., 2019; Wieczorek et al., 2020; Consolati, et al., 2020). In *Drosophila*, the composition of γTuRC during spermatogenesis differs from that at the centrosomal MTOC in other cells: Grip-84/GCP2, Grip-91/GCP3 and Grip128/GCP5 have testis-specific isotypes (Alzyoud et al., 2021), and Grip75/GCP4 and MOZART1 are specifically required for spermatogenesis (Vogt et al., 2006; Tovey et al., 2018). Thus, some components of γTuCs are linked specifically to the ncMTOC to control tissue-specific MT organization.

Because the composition and function of γTuC components vary in the ncMTOC of different tissues, the mechanism by which γTuC is recruited to the ncMTOC may also differ from that governing its recruitment to centrosomes. In embryonic intestinal epithelial cells in *C. elegans*, WDR-62 recruits γTuC to the apical ncMTOC, but WDR-62 is not involved in γTuC recruitment to centrosomes. On the other hand, MZT-1 is essential for centrosomal recruitment of γTuC but dispensable for its recruitment to the apical ncMTOC (Sallee et al., 2018; Sanchez et al., 2021). Thus, the composition and recruitment mechanism of γTuC appears to be diverse in mitotic centrosomes and ncMTOC in various tissues.

In the *C. elegans* germline, the mechanism by which γTuC is recruited to the ncMTOC on the plasma membrane remains unknown. Further studies of the tissue-specific roles and interactors of GTAP-1 and −2 will facilitate better understanding of the spatiotemporal regulation of ncMTOC positioning and function.

### Unique feature of MT dynamics in *C. elegans*

The unconventional composition of the *C. elegans* γTuC, i.e., with GTAP-1 and −2 instead of the typical GCP4–6, might correlate with certain unique features of MTs in this organism. First, *C. elegans* embryos have unusually high activity to assemble MTs in a γ-tubulin-independent manner. Whereas loss of γ-tubulin results in a drastic reduction of MTs in the majority of organisms, up to ~40% of MTs in *C. elegans* early embryos are assembled in a γ-tubulin-independent yet Aurora A protein 1–dependent manner (Hannak et al., 2002; Motegi et al., 2006; Toya et al., 2011). Electron microscopic analysis also revealed that *C. elegans* embryos have open-ended MTs and capped-ended MTs and that the latter are likely to be assembled with γTuC (O’Toole et al., 2003; O’Toole et al., 2012). One possibility is that an altered γTuC composition in ancestral nematodes of *C. elegans* might have reduced the ability to nucleate MTs, which led to the emergence of a compensatory γ-tubulin-independent (Aurora A protein 1–dependent) MT assembly mechanism. Second, the fact that the general number of MT protofilaments in most organisms is 13, corresponding to the 13-symmetrical structure of the γTuC, also has implications on the unconventional γTuC symmetry in *C. elegans*. The fundamental role of γTuC is to serve as a template for MTs, and a typical γTuC has 13-fold symmetry, which is consistent with the 13 protofilaments of canonical MTs in most eukaryotic cells. In contrast, *C. elegans* neurons and embryos generally have 11 profilaments (Chalfie and Thomson, 1982; Chaaban et al., 2018). It is tempting to speculate that the unconventional composition of the *C. elegans* γTuC, i.e., containing GTAP-1 and −2, may help restrict the number of MT protofilaments to 11 *in vivo*. Further structural studies will be needed to determine whether the *C. elegans* γTuC provides the structural seed to form 11-protofilament MTs.

## MATERIALS AND METHODS

### *C. elegans* strains

Strains of *C. elegans* were cultured using standard methods (Brenner, 1974) at 20°C (N2) or 24°C (all fluorescing strains). Table 1 lists the strains constructed in this study. Strain SA250 *[unc-119(ed3);tjIs54[pie-1p::gfp::tbb-2 pie-1p::2xmCherry::tbg-1 unc-119(+)]; tjIs57[pie-1p::mCherry:histoneH2 unc-119(+)]* (Toya et al., 2010) was used for the microscopic analysis by monitoring β-tubulin, γ-tubulin and histone.

**Table 1.**
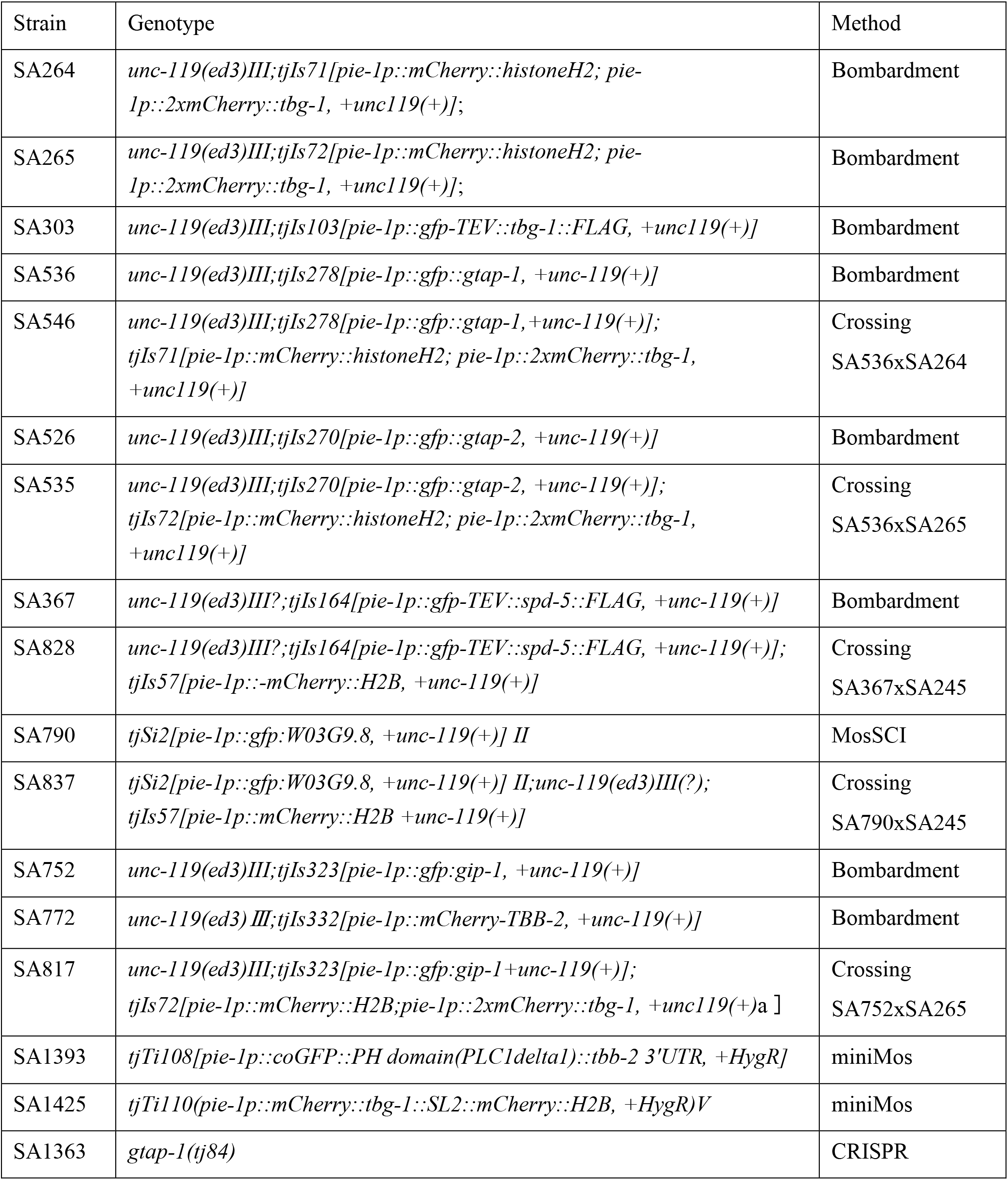
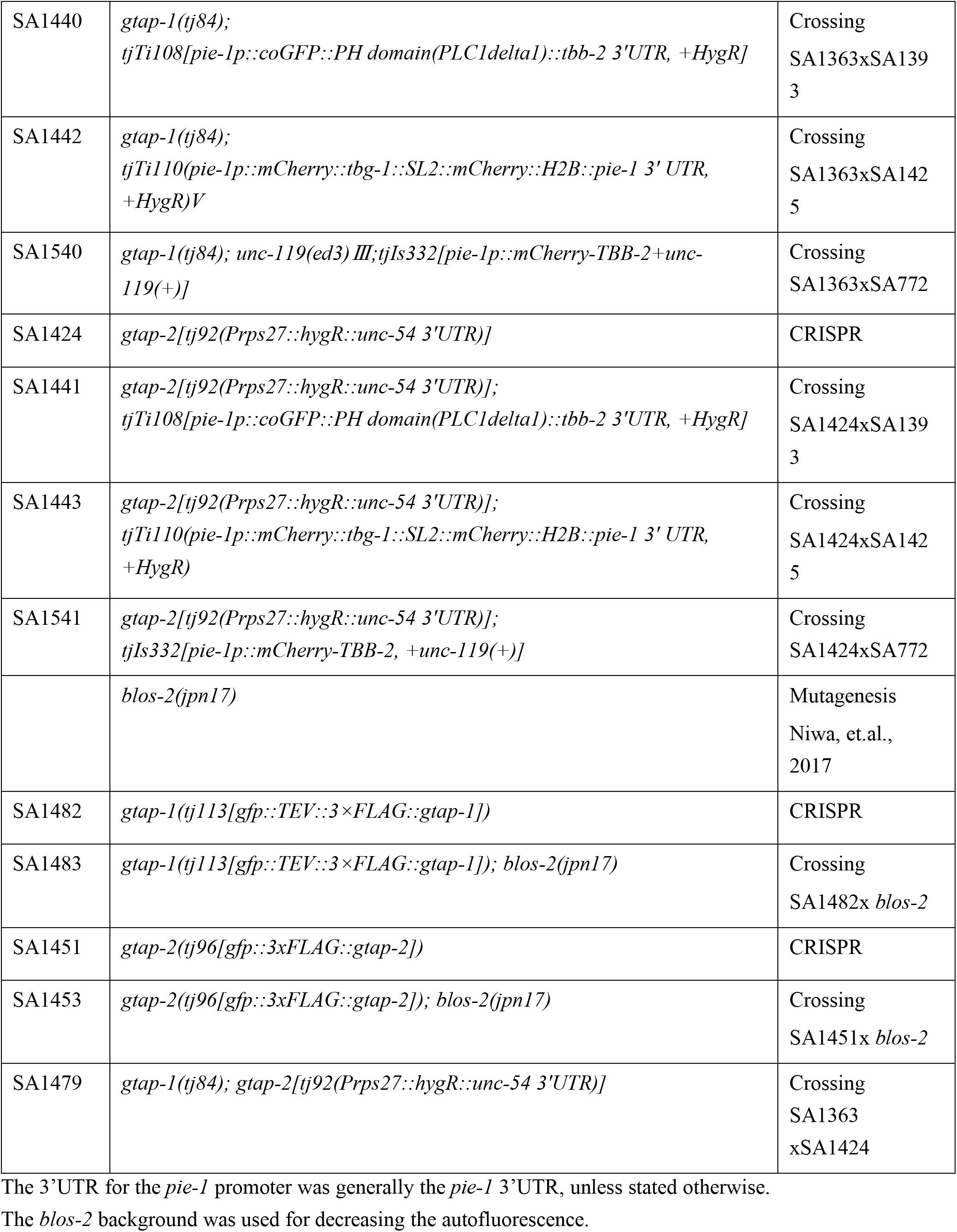
Strains constructed in this study.

The strains that expressed GFP::TEV::γ-tubulin::FLAG, GFP::GTAP-1, GFP::GTAP-2, GFP::SPD-5 and GFP::GIP-1 were constructed by high-pressure microparticle bombardment (Praitis et al., 2001) of DP38 [unc-119(ed3)] worms with plasmids pMTN1G_TEV::tbg-1(γ-tubulin)_FLAG, pMTN1G_gtap-1, pMTN1G_gtap-2, pMTN1G_spd-5, and pMTN1G_GIP-1, respectively. To construct these plasmids, full-length γ-tubulin with TEV and FLAG sequences, GTAP-1 and −2, SPD-5 and GIP-1 cDNAs was amplified by PCR and cloned into the entry vector pDONR201 (Thermo Fisher). Thereafter, the inserts were transferred to plasmid pMTN1G (Toya et al., 2010) by the LR reaction (Thermo Fisher).

To construct strain GFP::MZT-1 (W08G9.8), the MosSCI transposon method (www.wormbuilder.org, Frøkjær-Jensen et al., 2008) was used. In brief, the cDNA fragment encoding W08G9.8 was amplified and inserted into pCFJ350 (a gift from Erik Jorgensen, Addgene #34866) with the gene *gfp*, the *pie-1* promoter, and the *pie-1* 3’ untranslated region (UTR) by Gibson assembly (Gibson et al., 2009). The plasmid was injected into worms along with a plasmid containing transposase (pCFJ601, Peft-3::Mos1 transposase, Addgene #34874) and co-injection markers (pGH8 Addgene #19359, pCFJ90 Addgene #19327, pCFJ104 Addgene #19328, all gifts from Erik Jorgensen) into strain EG6699 [*ttTi5605 II; unc-119(ed3) III; oxEx1578*] to integrate into the ttTi5605 site.

To construct the strain that expressed GFP::PH and mCherry::TBG-1::SL2::mCherry::histone H2B (SA1425), we used the miniMos single-insertion method (www.wormbuilder.org, Frøkjær-Jensen et al., 2014). In brief, the PCR-amplified fragment of *pie-1*p::*gfp*::PH::*pie-1* 3’UTR was inserted into pCFJ1662 (a gift from Erik Jorgensen, Addgene #51482). Plasmid mixtures containing pCFJ1662_PH, pCFJ601, pMA122 (*peel-1*, Addgene #34873), and co-injection markers were injected into N2 young adult worms, and transgenic strains were obtained by hygromycin selection. To co-express the mCherry::TBG-1 (genomic DNA) and mCherry::histone H2B, plasmid pNH33 was generated by connecting two genes with a SL2 trans-splicing sequence and inserted into pCFJ910 (a gift from Erik Jorgensen, Addgene #44481) with the *pie-1p* and *pie-1* 3’UTR. Plasmid mixtures containing pNH33, pCFJ601, pMA122 and pBN41 (a gift from Peter Askjaer, Addgene #86716) were injected into N2 young adult worms, and transgenic strains were obtained by NeoR selection.

To construct the *gtap-1* null mutant, the CRISPR/Cas9 system was used with purified Cas9 proteins and single guide RNAs (sgRNAs) generated by *in vitro* transcription as described by Honda et al. (2017). In brief, Cas9 (2.5 μM final concentration) was injected with 10 μM sgRNA (target site: 5’-<u>CCT</u>AGAGATCCGTTTCCAGCTGT-3’) and 30 ng/μl ssODN (5’-GACATCTCACGTGCTGATTCTGCAGTATGTCTCACGACACGTT<u>GATATC</u>CGTTTCCAGCTGTTCTACAGGTAATTGAAGATGAACGAAAACTTTTTATA-3’), which introduced a stop codon (at the arginine 51 codon) and a cleavage site for EcoRV. For screening, two sgRNAs (2.5 μM each; 5’-CCGATGAGCATGGGATCCAGCCT-3’, 5’-<u>CCA</u>GCCTGATGGAACTTATAAGG-3’) were co-injected to introduce a mutation in *ben-1*.

To generate the *gtap-2* mutant, 1632 bp of the *gtap-2* coding region was substituted with the *Prps27::hygR::unc-54* 3’UTR fragment using the CRISPR/Cas9 system. To construct the template plasmid pNH32, ~500-bp homology arms were amplified that were adjacent to the double digestion sites within the *gtap-2* coding region. Plasmids for sgRNAs (pTK73_*gtap-2#3* and pTK73_*gtap-2#4*) were constructed by inserting the target sequences (5’-<u>CCC</u>GGTTGTCAGATGAGGATTT-3’ and 5’-TTCGATTCATTGTGAAGTAGGG-3’, respectively) into pTK73 as described by Obinata et al. (2018). Plasmids pNH32 (20 ng/μl), pTK73_*gtap-2#3*, pTK73_*gtap-2#4* (50 ng/μl), pDD162 (50 ng/μl, to express Cas9; a gift from Bob Goldstein, Addgene #47549) (Dickinson et al., 2013), and co-injection markers were injected and screened with 4 mg/ml hygromycin B. The endogenously GFP-tagged GTAP-1 and GTAP-2 strains were constructed as described by Dickinson et al. (2015). In brief, homology arms for *gtap-1* (upstream; 542 bp, downstream; 761 bp) and *gtap-2* (upstream; 588 bp, downstream; 668 bp) that had been amplified from the N2 genomic DNA were inserted into the ClaI-SpeI-cleaved pDD282 (a gift from Bob Goldstein, Addgene #66823, Dickinson et al., 2015) by Gibson assembly. To generate the endogenously GFP-tagged GTAP-1 strain, a plasmid mixture containing pDD162 (50 ng/μl), pDD282_*gtap-1* (50 ng/μl), and pTK73_*gtap-1_*F+52 (30 ng/μl) targeted to the sequence 5’-GGAATTCGATCAATGCACGC<u>AGG</u>-3’ and co-injection markers were injected. To generate the endogenously GFP-tagged GTAP-2 strain, a mixture containing Cas9 proteins (2.5 μM), pDD282_*gtap-2*#1 (50 ng/μl) and a synthesized sgRNA (2.5 μM; Integrated DNA Technologies) targeted to the sequence 5’-ATTAAGCTTTCTAATACATG<u>GGG</u>-3’ was injected. All strains were backcrossed with N2 and confirmed by DNA sequencing of the PCR-amplified modified region.

### Immunoprecipitation and mass spectrometry

Young adult worms expressing GFP::TBG-1::FLAG (SA303) were grown synchronously on EPP plates (25mM potassium phosphate (pH 6.0), 20 mM NaCl, 2 % (w/v) peptone, 1mM MgSO4, 5 μg/ml cholesterol, 2.5 % (w/v) agar). Approximately 3 million embryos were collected by bleaching (Epstein and Shakes, 1995), washed in lysis buffer (50 mM HEPES pH 7.4, 1 mM EGTA, 1 mM MgCl2, 100 mM KCl, 10% (v/v) glycerol, 0.05% (v/v) NP40, and 0.1 mM GTP), and frozen in liquid nitrogen. Embryos were suspended in 1 ml lysis buffer containing a protease inhibitor cocktail (Roche) supplemented with 1 mM phenylmethylsulfonyl fluoride and lysed by sonication. After centrifugation at 20,000 × *g* for 10 min, 20 μg anti-FLAG preincubated with 50 μl protein G–conjugated Dynabeads (Thermo Fisher) was mixed with the supernatant for 3 h at 4°C. Immunoprecipitates were collected using magnets, washed three times with 1 ml lysis buffer, and immunoprecipitated proteins were eluted from Dynabeads by incubating with 200 μM FLAG peptides (Sigma-Aldrich) for 90 min. Elutes were subjected to precipitation with acetone, lysed in SDS-PAGE loading buffer, subjected to gradient SDS-PAGE (5–20% polyacrylamide), and visualized by silver staining. Bands specifically detected in the extract of strain SA303 were excised from the gel and analyzed by mass spectrometry. For a comprehensive analysis, each lane was dissected into 15 pieces using a razor blade, and each gel piece was analyzed by mass spectrometry.

The number of peptides for each protein detected by mass spectrometry was compared between the GFP::γ-tubulin::FLAG-expressing and control strains. Proteins with two or more identified peptides were selected to ensure the reliability of the results. Thereafter, proteins were selected if they were enriched by ≥3-fold in strain SA303 compared with the control strain. Finally, proteins that were abundant in embryos and unlikely to have a functional relationship with γ-tubulin (e.g., vitellogenin) were excluded. The remaining proteins were considered as potential GFP::γ-tubulin::FLAG interactors.

### Antibodies

To raise rabbit polyclonal antibodies specific for GIP-1 and GTAP-1, a cDNA fragment corresponding to amino-acid residues 771–891 of GIP-1 **or** the full-length GTAP-1 cDNA were inserted into pDEST17 (Thermo Fisher). The resultant fusion proteins were expressed in *Escherichia coli*, purified using nickel affinity columns, and used to inoculate rabbits (carried out by Medical and Biological Laboratories). To generate anti-GIP-2, full-length GIP-2 cDNA with a C-terminal V5-His6 tag was inserted into pColdI (TAKARA) and used to inoculate rabbits and rats. Antibodies to GIP-1, −GIP-2, and −GTAP-1 were affinity purified. Rabbit anti-GTAP-2 was generated by inserting residues 591–823 of GTAP-2 into vector pColdI. Serum from GTAP-2-immunized rabbits was fractionated to obtain a IgG-enriched fraction. Rat anti-γ-tubulin was prepared as described by Toya et al. (2011). The secondary antibodies for western blotting were horseradish peroxidase–conjugated anti-rabbit and horseradish peroxidase–conjugated anti-rat (Jackson ImmunoResearch).

### Yeast two-hybrid analysis

Yeast two-hybrid analysis was performed using the ProQuest Two-Hybrid system (Thermo Fisher). Briefly, PCR-amplified cDNA fragments encoding each protein were subcloned in-frame downstream of the GAL4 DNA-binding domain of pDEST32 or the GAL4 activation domain of pDEST22. The constructed bait and prey vectors were confirmed by DNA sequencing.

For expression of γ-tubulin/TBG-1 or GIP-2/GCP3 as the “third” protein, i.e., in addition to the DNA-binding domain–fused and activation domain–fused proteins, the *ADH1* promoter was inserted at the HindIII-SmaI site in plasmid pRS426, which contains the URA3 gene; then, the TBG-1 or GIP-2 coding sequences were inserted downstream of the *ADH1* promoter. For expression of γ-tubulin/TBG-1 or MZT-1 as a “fourth” protein, pAUR112 (TAKARA) carrying the Aureobasidin A resistance gene was digested with BstBI and SmaI to delete the URA3 gene, and then the *ADH1* promoter and the TBG-1 or MZT-1 coding sequence were inserted into the KpnI-SacI site.

For typical yeast two-hybrid assays, yeast strain Mav203 was co-transformed with a bait vector and a prey vector and incubated on Sc-LeuTrp plates. For each pair of vectors transformed, four transformants were picked and characterized on selection plates for 3–4 days at 30°C.

For expression with γ-tubulin/TBG-1 as the “third” protein, Mav203 was transformed with three plasmids and spread onto Sc-LeuTrpUra plates. For co-expression with the “third” and “fourth” proteins, Mav203 was transformed with four plasmids and spread onto Sc-LeuTrpUra plates containing 0.5 μg/ml Aureobasidin A (TAKARA).

### Sucrose gradient sedimentation

*Caenorhabditis elegans* eggs (100 μl, ~30,000/μl, stored at –80°C) were collected by bleaching and suspended in 300 μl lysis buffer containing 50 mM HEPES pH 7.5, 1 mM MgCl_2_, 1 mM EGTA, 1 mM β-mercaptoethanol, 100 mM NaCl, 0.1 mM GTP, 1 mM phenylmethylsulfonyl fluoride, and protease inhibitors (Roche). A crude extract was prepared by sonicating the eggs six times for 10 s each, followed by centrifugation at 20,000 × *g* for 15 min at 4°C. The supernatant (200 μl) was loaded onto a 3.8-ml, 4–40% sucrose gradient prepared in lysis buffer and centrifuged at 150,000 × *g* for 4 or 16 h at 4°C in a MLS-50 rotor (Beckman Coulter). Thereafter, 20 fractions (200 μl) were collected and analyzed by western blotting using affinity-purified rat anti-γ-tubulin (1:1000), rabbit anti-GIP-1/GCP3 (1:1000), rabbit anti-GTAP-1 (1:300), and rat anti-GFP (1:1000). GIP-2/GCP2 was probed using anti-GIP-2/GCP2 serum (1:1000).

### RNAi

RNAi was carried out using the soaking method. The following cDNA clones (gifts from Professor Yuji Kohara, National Institute of Genetics, Mishima, Japan) were used as templates to synthesize double-stranded RNAs (dsRNAs): yk1562g08 (*tbg-1*), yk330f6 (*gtap-1*), and yk1443f05 (*gtap-2*). The cDNA inserts were PCR-amplified using primer sets containing vector and T7 primer sequences. Primers Cmo422 and T7 were used to amplify *gtap-1*, whereas primers T7-ME774 and T7-ME1250 were used to amplify *tbg-1* and *gtap-2*, respectively. The primer sequences were as follows: Cmo422, 5′-GCGTAATACGACTCACTATAGGGAACAAAAGCTGGAGCT-3′; T7, 5′-GTAATACGACTCACTATAGGGC-3′; T7-ME774, 5′-TAATACGACTCACTATAGGGCTTCTGCTCTAAAAGCTGCG-3′; and T7-ME1250, 5′-TAATACGACTCACTATAGGGTGTGGGAGGTTTTTTCTCTA-3′. For *mzt-1* and *gip-1*, the cDNA templates were amplified from a cDNA library (Invitrogen) using gene-specific primers containing the T7 promoter sequence at each 5’ end. Primers for *mzt-1* were T7_mzt-1_F (5’-TAATACGACTCACTATAGGGATGAGCGACCCAAAGAAACAC-3’) and T7_mzt-1_R (5’-TAATACGACTCACTATAGGGTCATGACAATGCATTTTCCCG-3’), and primers for *gip-1* were T7_gip-1_F(5’-TAATACGACTCACTATAGGGATGCGTCGACAAGGCAGCGAA-3’) and T7_gip-1_R_270AA (5’-TAATACGACTCACTATAGGGCACAGATGTATTCAATAGGTGG-3’). Then, dsRNAs were synthesized *in vitro* using the RiboMAX T7 Express System (Promega) and purified by the phenol chloride.

Worms were soaked in 2 mg/ml dsRNA solution at 24°C for 12–24 h. After removal from the dsRNA solution, worms were cultured at 24.5°C for 24 h, then observed. For partial RNAi of *tbg-1*, worms were soaked in 0.3 mg/ml *tbg-1* dsRNA solution at 24.5°C for 2 h. After removal from the dsRNA solution, worms were cultured at 24.5°C for 10–18.5 h and observed. The knockdown efficiencies of GTAP-1 and −2 were confirmed by staining RNAi-treated embryos with an antibody specific for each target protein.

### Microscopic imaging of embryos

For time-lapse microscopy, embryos expressing fluorescent proteins were mounted on 2% agarose pads. The specimens were imaged using a CSU-X1 spinning-disc confocal system (Yokogawa Electric Corp.) mounted on an IX71 inverted microscope (Olympus) controlled by MetaMorph software (Molecular Devices). Images were acquired as described by Toya et al. (2010). Briefly, images were acquired using an Orca-R2 12-bit/16-bit cooled charge-coupled device camera (Hamamatsu Photonics) and a 60× 1.30 NA UPlanSApo silicon objective lens without binning and with streaming. To obtain Z-sectioned images, 7–25 Z sections at 1-μm steps were acquired using a 300- to 500-ms exposure (camera gain, 255) for each wavelength. For time-lapse recording, images were acquired every 30 s. To analyze colocalization, three Z-sectioned images of GFP and mCherry at each Z position were obtained at an interval of 30 s without streaming. To assess the dependency of GTAP-1 and −2 localization on TBG-1/γ-tubulin, 25 Z-sectioned images were obtained for *tbg-1(RNAi)* embryos to cover the Z axis of the entire embryo, and single Z-sectioned images of centrosomes were obtained in control embryos to avoid a time lag between the 488-nm and 568-nm excitation.

To quantify the fluorescence intensities of TBG-1, GTAP-1, and GTAP-2 in the centrosomal region, projected images of seven Z-sectioned images with a Z interval of 1 μm around centrosomes were generated using the SUM algorithm in ImageJ software (National Institutes of Health). The average intensity was measured in a circular region of radius 3 μm around each centrosome in the projected images, and the average fluorescence intensity in a cytoplasmic ring-shaped region around centrosomes was subtracted. The cytoplasmic region had an inner radius of 3 μm and an outer radius of 4.2 μm in one-cell embryos. To quantify MTs, the average fluorescence intensity was measured in a circular region of radius 6 μm around each centrosome, and the average fluorescence in a circular region of radius 6 μm outside embryos was subtracted as background. Fluorescence intensities were normalized by the average fluorescence intensity in control embryos. Projection and quantification were performed using ImageJ/Fiji software. Statistical analysis was performed using GraphPad Prism software (MDF Co.).

### Microscopic imaging of the adult germline

For live-cell imaging of the GFP::PH domain and **e**ndogenously tagged GFP::GTAP-1 and GFP::GTAP-2 in the adult gonads, adult worms were treated with 0.0025 mM levamisole in polystyrene polybeads (Polysciences) and mounted on 5% agar pads. For imaging, the same microscopic instruments were used as described above with a 60× 1.3 NA UPlanSApo silicon objective lens for the GFP::pH domain and mCherry::tbb-2, and a 100× 1.35 NA UPlansSApo silicon objective lens for endogenously tagged GFP::GTAP-1 and GFP::GTAP-2. Image projection and tiling were performed using NIH ImageJ/Fiji software.

For imaging mCherry::histone and γ-tubulin in adult gonads, the same microscopic instruments were used as described above with a 60× 1.3 NA UPlanSApo silicon objective lens except that the camera used was ORCA-Flash 4.0 sCMOS (Hamamatsu Photonics). To quantify the amount of γ-tubulin at the plasma membrane, the fluorescence intensities in the band regions (0.85 × 8.5 μm) were measured using Prot Profile in Fiji software, and the mCherry::γ-tubulin signals across the plasma membrane were normalized using that of mCherry::histone.

### Phylogenetic analysis

Orthologs of tubulins and the components of γTuC were searched by BLAST (Altschul, et al., 1990), PSI-BLAST (position-specific iterated BLAST) (Altschul, et al., 1997), and DELTA-BLAST (Domain Enhanced Lookup Time Accelerated BLAST) (Boratyn, et al., 2012) against human orthologs. The sequence similarity between orthologs was calculated and phylogenetic trees were drawn using ClustalW (EMBL-EBI).

## ACKNOWLEDGEMENTS

We are grateful to Professors Masakado Kawata and Takashi Makino (Tohoku University, Graduate School of Life Science) for discussion pertaining to molecular evolution, Professor Yuji Kohara (National Institute of Genetics, Mishima, Japan) for providing cDNA clones, and Ms. Hiroko Sugawara, Ms. Makiko Sasaki, Dr. Kenji Tsuyama, Dr. Satoshi Namai and Ms. Yuki Hoshi for helping construct the plasmids and worm strains. Some strains were provided by the Caenorhabditis Genetics Center, which is funded by the NIH Office of Research Infrastructure Programs (P40 OD010440). This work was supported by JSPS KAKENHI Grant Numbers JP15H04369, JP15K14503 and Bilateral Joint Research Project (to AS) and JP16K07334 and JP20K06616 (to NH). This work was partially supported by a Tohoku University Center for Gender Equality Promotion (TUMUG) Support Project and by grants-in-aid and the Toyota Riken Scholar from the Toyota Physical and Chemical Research Institute (to NH).

## AUTHOR CONTRIBUTIONS

Conceptualization: N. Haruta, A. Sugimoto; Investigation: N. Haruta, E. Sumiyoshi, Y, Honda, M. Terasawa, M. Toya, C. Uchiyama; Writing - original draft: N. Haruta, E. Sumiyoshi, Y. Honda; Writing - review & editing: N. Haruta, A. Sugimoto; Project administration: A. Sugimoto; Funding acquisition: N. Haruta, A. Sugimoto

---

**Supplemental Fig. 1.**
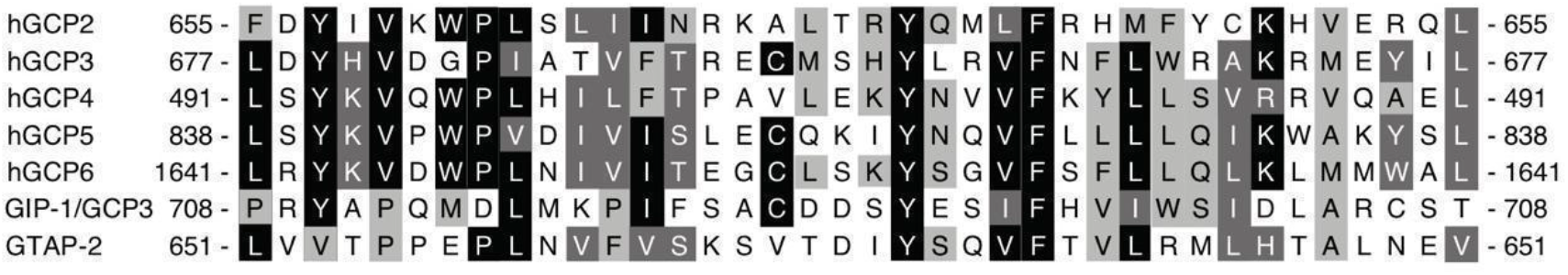
Amino-acid sequence alignment of partial regions of the grip-2 motif in GTAP-2, GIP-1/GCP3, and human GCPs (hGCP2~6). The extent of conservation is indicated by shading (black = greatest conservation, light gray = least).

**Supplemental Fig. 2.**
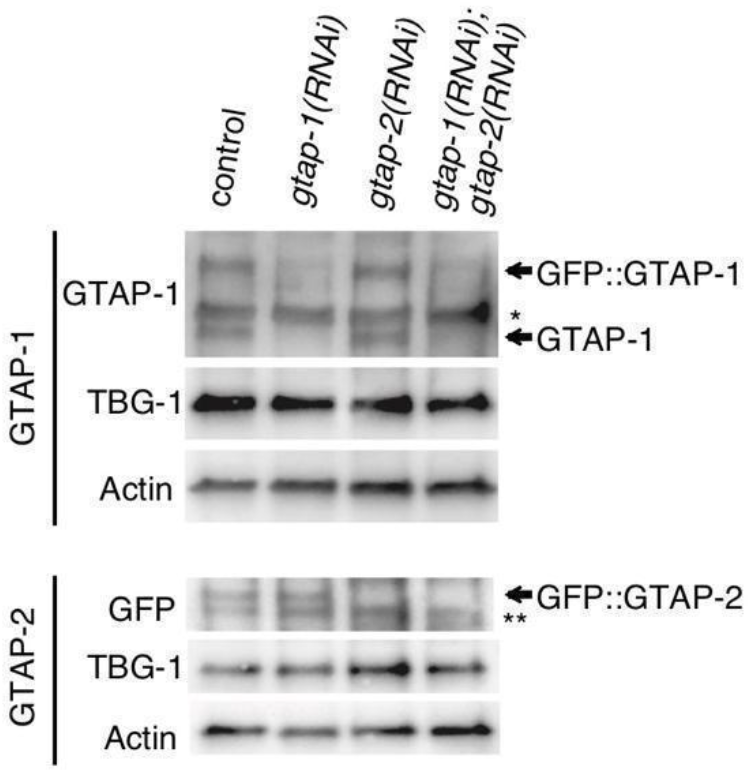
Western blot analysis of RNAi-mediated knockdown in GFP::GTAP-1- and GFP::GTAP-2-expressing strains. Anti-GTAP-1 was used to detect GTAP-1 and GFP::GTAP-1. Anti-GFP was used to detect GFP::GTAP-2. The intensities of the bands corresponding to GFP::GTAP-1 and GTAP-1 were significantly lower in *gtap-1(RNAi)* and *gtap-1(RNAi);gtap-2(RNAi)* worms than in the control and *gtap-2(RNAi)* worms. The intensities of the bands corresponding to GFP::GTAP-2 were significantly lower in *gtap-2(RNAi)* and *gtap-1(RNAi);gtap-2(RNAi)* worms than in the control and *gtap-1(RNAi)* worms.

**Supplemental Fig. 3.**
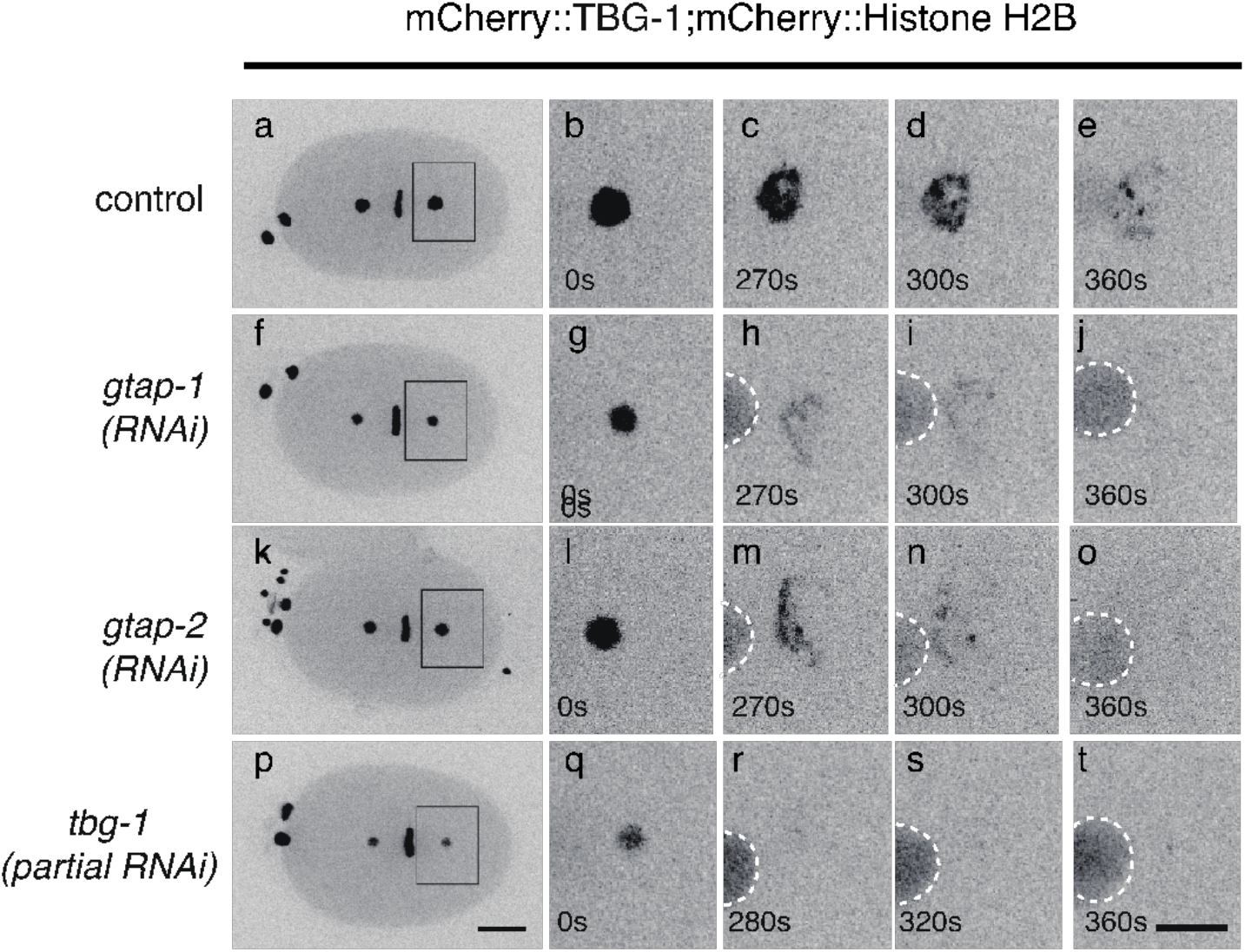
GTAP-1 and GTAP-2 affect γ-tubulin dynamics during mitosis. Time-lapse confocal micrographs of mCherry::γ-tubulin and mCherry::histone in the control, *gtap-1(RNAi)*, *gtap-2(RNAi)*, and *tbg-1(partial RNAi)* embryos. The far-left panels show images of whole embryos. The right panels are magnified views of the centrosomal regions in time-lapse images. Times are relative to metaphase. White dotted lines indicate nuclei. Scale bars, 10 μm (a, f, k, and p) or 5 μm (b–e, g–j, l–o, and q–t).

**Supplemental Fig. 4.**
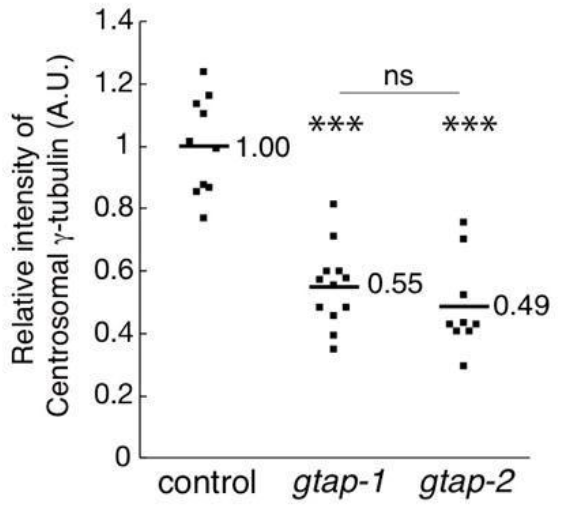
Fluorescence intensities of mCherry::γ-tubulin at the centrosome in the control, *gtap-1(RNAi)*, and *gtap-2(RNAi)*. Statistical analysis was carried out with the Tukey-Kramer multiple comparison test for three groups; \*\*\**p*<0.0001.

**Supplementary Table 1.**
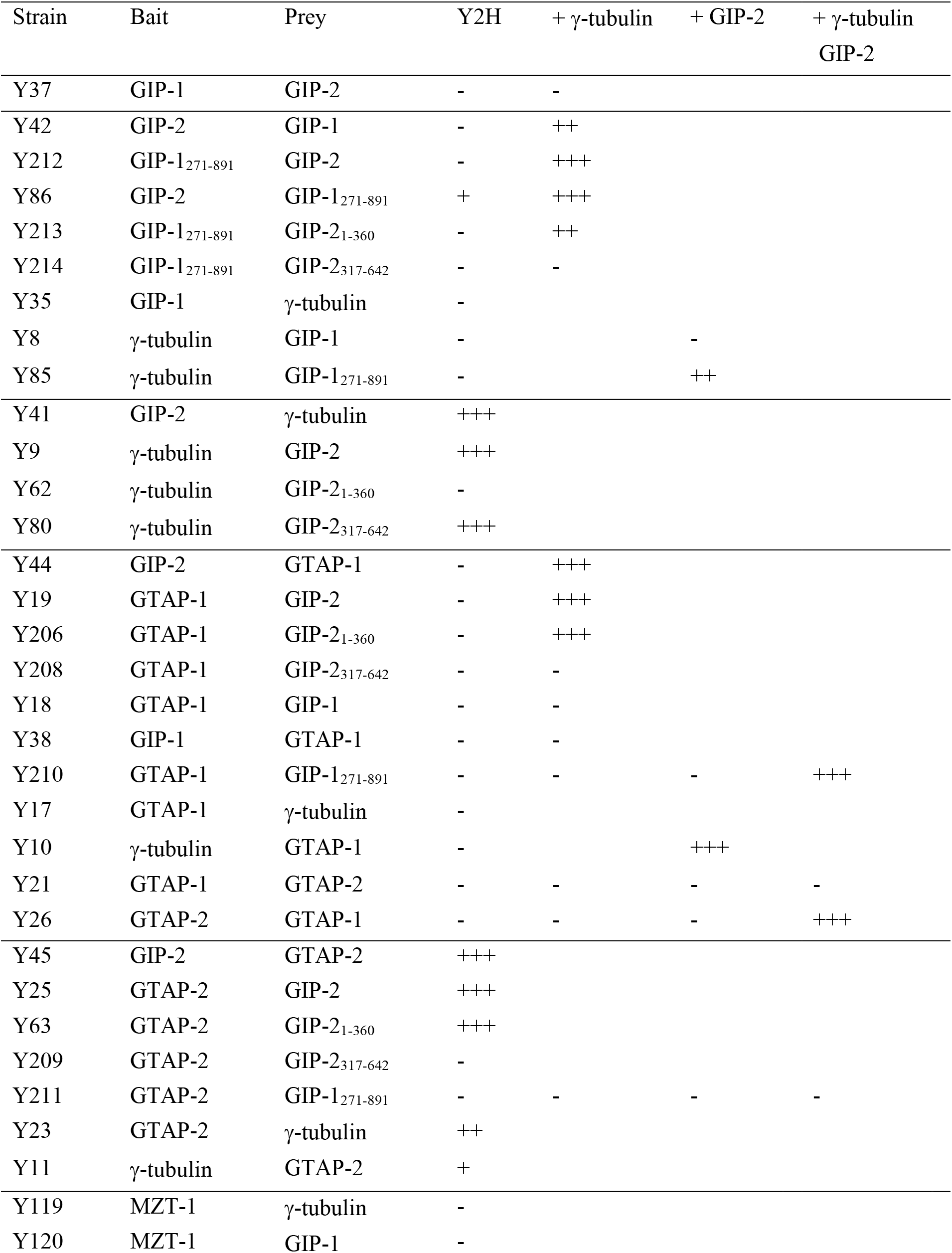
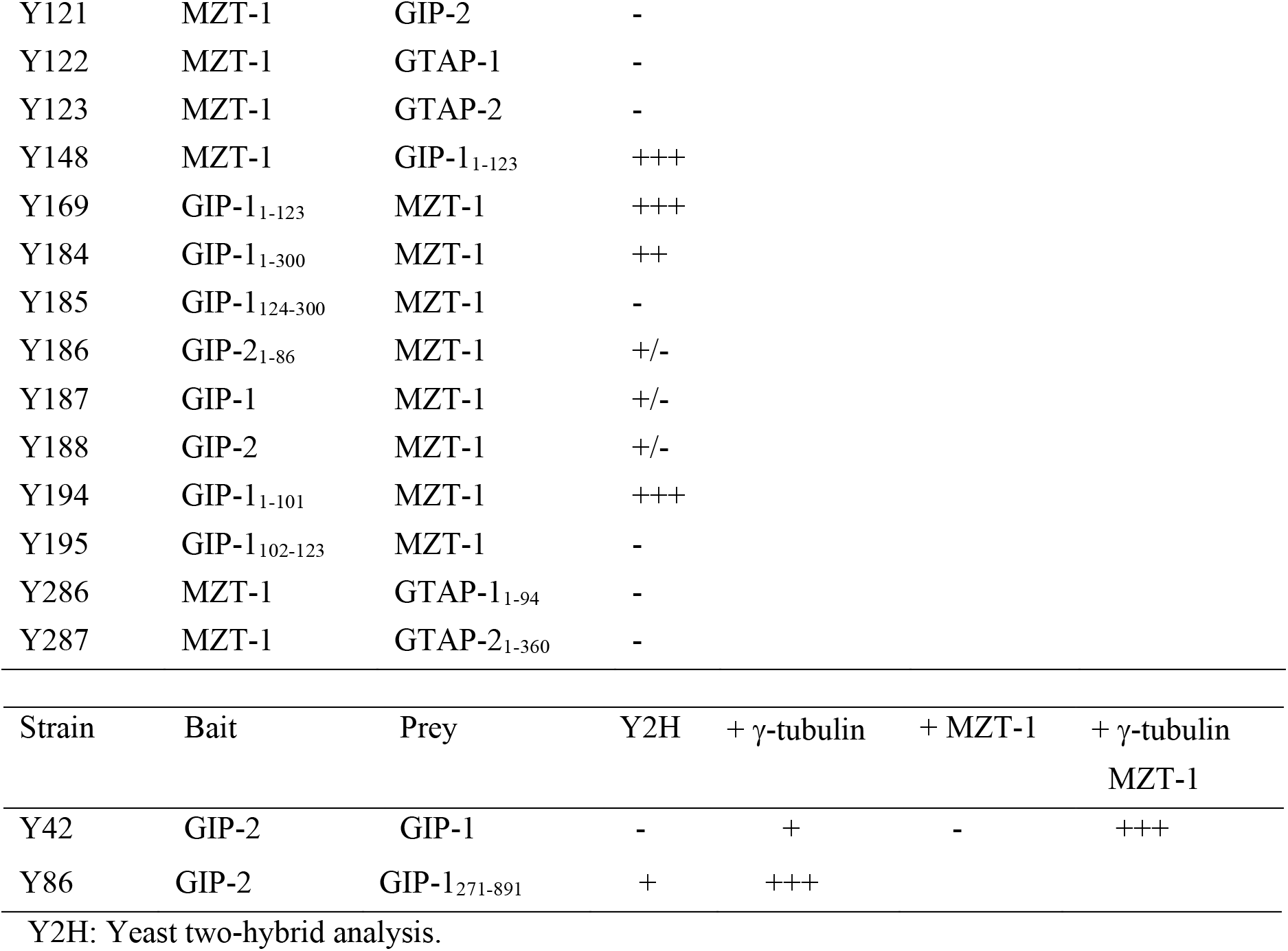
Interactions identified by yeast two-hybrid assays with and without γ-tubulin, GIP-2/GCP2, and/or MZT-1.

| Strain | Bait | Prey | Y2H | + γ-tubulin | + GIP-2 | + γ-tubulin<br>GIP-2 |
| --- | --- | --- | --- | --- | --- | --- |
| Y37 | GIP-1 | GIP-2 | - | - |  |  |
| Y42 | GIP-2 | GIP-1 | - | ++ |  |  |
| Y212 | GIP-1<sub>271-891</sub> | GIP-2 | - | +++ |  |  |
| Y86 | GIP-2 | GIP-1<sub>271-891</sub> | + | +++ |  |  |
| Y213 | GIP-1<sub>271-891</sub> | GIP-2<sub>1-360</sub> | - | ++ |  |  |
| Y214 | GIP-1<sub>271-891</sub> | GIP-2<sub>317-642</sub> | - | - |  |  |
| Y35 | GIP-1 | γ-tubulin | - | | | |
| Y8 | γ-tubulin | GIP-1 | - | | - | |
| Y85 | γ-tubulin | GIP-1<sub>271-891</sub> | - | | ++ | |
| Y41 | GIP-2 | γ-tubulin | +++ | | | |
| Y9 | γ-tubulin | GIP-2 | +++ | | | |
| Y62 | γ-tubulin | GIP-2<sub>1-360</sub> | - | | | |
| Y80 | γ-tubulin | GIP-2<sub>317-642</sub> | +++ | | | |
| Y44 | GIP-2 | GTAP-1 | - | +++ |  |  |
| Y19 | GTAP-1 | GIP-2 | - | +++ |  |  |
| Y206 | GTAP-1 | GIP-2<sub>1-360</sub> | - | +++ |  |  |
| Y208 | GTAP-1 | GIP-2<sub>317-642</sub> | - | - |  |  |
| Y18 | GTAP-1 | GIP-1 | - | - |  |  |
| Y38 | GIP-1 | GTAP-1 | - | - |  |  |
| Y210 | GTAP-1 | GIP-1<sub>271-891</sub> | - | - | - | +++ |
| Y17 | GTAP-1 | γ-tubulin | - | | | |
| Y10 | γ-tubulin | GTAP-1 | - | | +++ | |
| Y21 | GTAP-1 | GTAP-2 | - | - | - | - |
| Y26 | GTAP-2 | GTAP-1 | - | - | - | +++ |
| Y45 | GIP-2 | GTAP-2 | +++ |  |  |  |
| Y25 | GTAP-2 | GIP-2 | +++ |  |  |  |
| Y63 | GTAP-2 | GIP-2<sub>1-360</sub> | +++ |  |  |  |
| Y209 | GTAP-2 | GIP-2<sub>317-642</sub> | - |  |  |  |
| Y211 | GTAP-2 | GIP-1<sub>271-891</sub> | - | - | - | - |
| Y23 | GTAP-2 | γ-tubulin | ++ | | | |
| Y11 | γ-tubulin | GTAP-2 | + | | | |
| Y119 | MZT-1 | γ-tubulin | - | | | |
| Y120 | MZT-1 | GIP-1 | - |  |  |  |

|  |  |  |  |
| --- | --- | --- | --- |
| Y121 | MZT-1 | GIP-2 | - |
| Y122 | MZT-1 | GTAP-1 | - |
| Y123 | MZT-1 | GTAP-2 | - |
| Y148 | MZT-1 | GIP-1<sub>1-123</sub> | +++ |
| Y169 | GIP-1<sub>1-123</sub> | MZT-1 | +++ |
| Y184 | GIP-1<sub>1-300</sub> | MZT-1 | ++ |
| Y185 | GIP-1<sub>124-300</sub> | MZT-1 | - |
| Y186 | GIP-2<sub>1-86</sub> | MZT-1 | +/- |
| Y187 | GIP-1 | MZT-1 | +/- |
| Y188 | GIP-2 | MZT-1 | +/- |
| Y194 | GIP-1<sub>1-101</sub> | MZT-1 | +++ |
| Y195 | GIP-1<sub>102-123</sub> | MZT-1 | - |
| Y286 | MZT-1 | GTAP-1<sub>1-94</sub> | - |
| Y287 | MZT-1 | GTAP-2<sub>1-360</sub> | - |

| Strain | Bait | Prey | Y2H | + γ-tubulin | + MZT-1 | + γ-tubulin<br>MZT-1 |
| --- | --- | --- | --- | --- | --- | --- |
| Y42 | GIP-2 | GIP-1 | - | + | - | +++ |
| Y86 | GIP-2 | GIP-1<sub>271-891</sub> | + | +++ |  |  |

Y2H: Yeast two-hybrid analysis.

